# Predicting recovery trajectories and injury severity following partial crush spinal cord injury in mice

**DOI:** 10.64898/2026.02.28.708735

**Authors:** Kunyu Li, Laboni F. Hassan, Himagowri Prasad, Glory C. Omodia, Paige S. Woods, Timothy M. O’Shea

## Abstract

The partial crush spinal cord injury (SCI) model enables preclinical testing of experimental therapies in mice, but substantial inter-animal variability in recovery outcomes confounds efficacy assessments. Here, we used open field behavioral data collected during the first 3 days post partial thoracic SCI to generate an Acute Functional Score (AFS) that defined three subgroups with divergent recovery trajectories. Applying latent class growth analysis and growth mixture modeling to open field and grid walk testing data, we demonstrated 83-92% prediction accuracy for AFS-defined recovery trajectories. The three subgroups differed significantly in treadmill kinematics and histological assessments of lesion size and astrocyte bridging. Applying the recovery trajectory framework to mice receiving saline or biomaterial vehicle injections at 3 days post-SCI revealed robust predictive accuracy while exposing disproportionate injury severity distributions between experimental groups. The approach enables individualized post-SCI recovery characterization that can neutralize procedural bias, minimize animal numbers, and provide a probabilistic basis for evaluating whether interventions enhance or suppress wound repair processes. Our findings establish a foundation for improving preclinical SCI study design and accelerating identification of effective therapies.

## Introduction

The partial crush spinal cord injury (SCI) paradigm is a popular model in pre-clinical rodent research that involves using forceps with a precise closure spacing to apply a controlled compression injury to the thoracic spinal cord^1^. This model induces an incomplete SCI by sparing portions of neural tissue within the affected segment, allowing for a natural course of spontaneous recovery that correlates with injury severity and that can be modulated by experimental treatments^1,2^. The pathophysiology of forcep crush resembles that of common clinical human SCI, making it a relevant model for studying the fundamental biology of wound healing after SCI^2–4^, and evaluating the efficacy of therapeutic strategies such as stimulated axon regeneration^5,6^ and neural cell grafting^7^. However, much like impactor contusion models^8^, the partial crush technique is associated with substantial inter-animal variability in injury severity, even when performed by experienced rodent surgeons. This variability is conferred during the crush surgery but is often not identified until the final histological analysis, if at all. Consequently, inconsistent injury severities can result in divergent spontaneous recovery outcomes on behavioral tests, allowing any procedural bias to significantly mask or exaggerate the efficacy of experimental interventions^9,10^. Detecting treatment effects in the presence of such variability necessitates highly powered studies with large sample sizes, which entail increased costs and resource demands, while also posing ethical concerns. To address these challenges, adopting analytical approaches that assess treatment effects on a subgroup or per animal basis, rather than relying solely on population-level statistical analyses, may improve sensitivity and reduce bias, thereby enhancing our ability to identify new effective therapies for SCI.

We are using the partial crush SCI model to test new therapeutic strategies to enhance glia-based wound repair. As part of these efforts, experimental interventions are surgically delivered locally during the sub-acute injury phase (1-5 days post injury) to modulate endogenous wound repair processes. Given the inherent experimental variability of the partial crush model, an analytical framework capable of distinguishing treatment-induced effects on functional recovery from those arising from natural spontaneous recovery is essential. To achieve this, a methodology for predicting the natural course of recovery from sparse information available only prior to application of treatment is needed. In clinical human SCI, there have been significant advancements in using acute injury data to predict long-term recovery outcomes^11–15^. Machine learning approaches have been applied to large prospective databases, such as the European Multicenter Study about Spinal Cord Injury (EMSCI), to model the spontaneous recovery trajectories of patients^11–15^. These studies have incorporated data from neurological impairment scales (e.g. ASIA) in conjunction with acute MRI imaging^16^, hematological information^17^, proteomic analysis^18^, surgery management information, and pertinent patient demographics. Often these predictor variables are limited to within the first several days after injury^19^. Recent clinical trials of experimental therapies in early stages of investigation have employed historical SCI conversion data as a part of their treatment effect sizes estimates in lieu of running a parallel placebo control group^20^. Despite these clinical advances, comparable predictive frameworks remain underdeveloped and seldom integrated into preclinical SCI research^21,22^.

Here, we present a data-driven analytic framework to predict the trajectory of natural functional recovery in mice subjected to the standardized partial crush SCI employed by our lab. The framework uses data collected from non-parametric open field locomotion testing during the acute injury period (days 1 to 3 post-SCI) and employs statistical methods, including latent class growth analysis and growth mixture modeling, to classify a large population of partial crush SCI mice training data into 3 distinct recovery subgroups. These classified subgroups show divergent recovery outcomes in open field locomotion, grid walk testing, and Treadmill based hindlimb kinematics analysis as well as significant differences in lesion sizes by post-hoc histological assessments. When applied to a separate validation cohort of mice receiving vehicle injections of saline or an inert biomaterial carrier, the subgroup classification model demonstrated strong predictive accuracy and robustness, enabling it to identify and neutralize the impact of inadvertent procedural bias. This analytical framework allows individualized characterization of recovery trajectories in mice after partial crush SCI, providing a probabilistic basis for evaluating whether a therapeutic intervention enhances or suppresses the expected recovery course, as well as a decision-support tool for optimizing preclinical study design.

## Results

### Partial crush SCI model shows highly variable natural post injury motor recovery

We and others employ a thoracic partial crush SCI model in mice, performed under general anesthesia by first exposing the spinal cord through a laminectomy at the T10 vertebra and then inducing a laterally applied crush injury at the corresponding T12 spinal segment^23^. We have modified a pair of McPherson straight tying forceps typically employed in clinical ophthalmic applications to fabricate our partial crush forcep tool (**Figure 1a**). These forceps are readily commercially available and contain a fenestrated handle which we leveraged to introduce a spacer to create a defined closure gap by inserting an 8mm silver grommet into one of the grip spacers using a hand press. The forceps without any modifications had a tip width of 0.5mm and the inserted grommet was adapted to achieve a consistent spacing upon closure of approximately 400µm (Mean ± SD = 401±13µm, Normally distributed) (**Figure 1b,c**), which we know from previous work by us and others should confer a moderately severe SCI in adult mice^1,3,4^. The inclusion of the grommet on the forcep provided a defined location for the surgeon to place their thumb during the crush procedure, ensuring consistent force was applied at the same fulcrum point on the forcep arm and creating reproducible, precise spacing upon closure. After subjecting the exposed spinal cord to the forcep crush, there is notable evidence of hemorrhage in deep spinal cord tissue regions when viewed from the dorsal surface of the cord, but the dorsal spinal vein (dSV) typically remains intact (**Figure 1d**). The extent of injury severity in these partial crush lesions is less obvious when compared to the distinct banded injury zone observed in the complete crush injury paradigm. We collected data from partial SCI crush surgeries performed on female and male wildtype C57BL/6 mice using this spaced forcep tool over a two-year period across 3 different surgeons, with these mice generated as part of surgical training of new graduate students and lab start-up pilot experiments. These accrued untreated “control” mice formed the basis of training data for the exploration of the predictive modeling of natural recovery trajectories outlined here.

**Figure 1:**
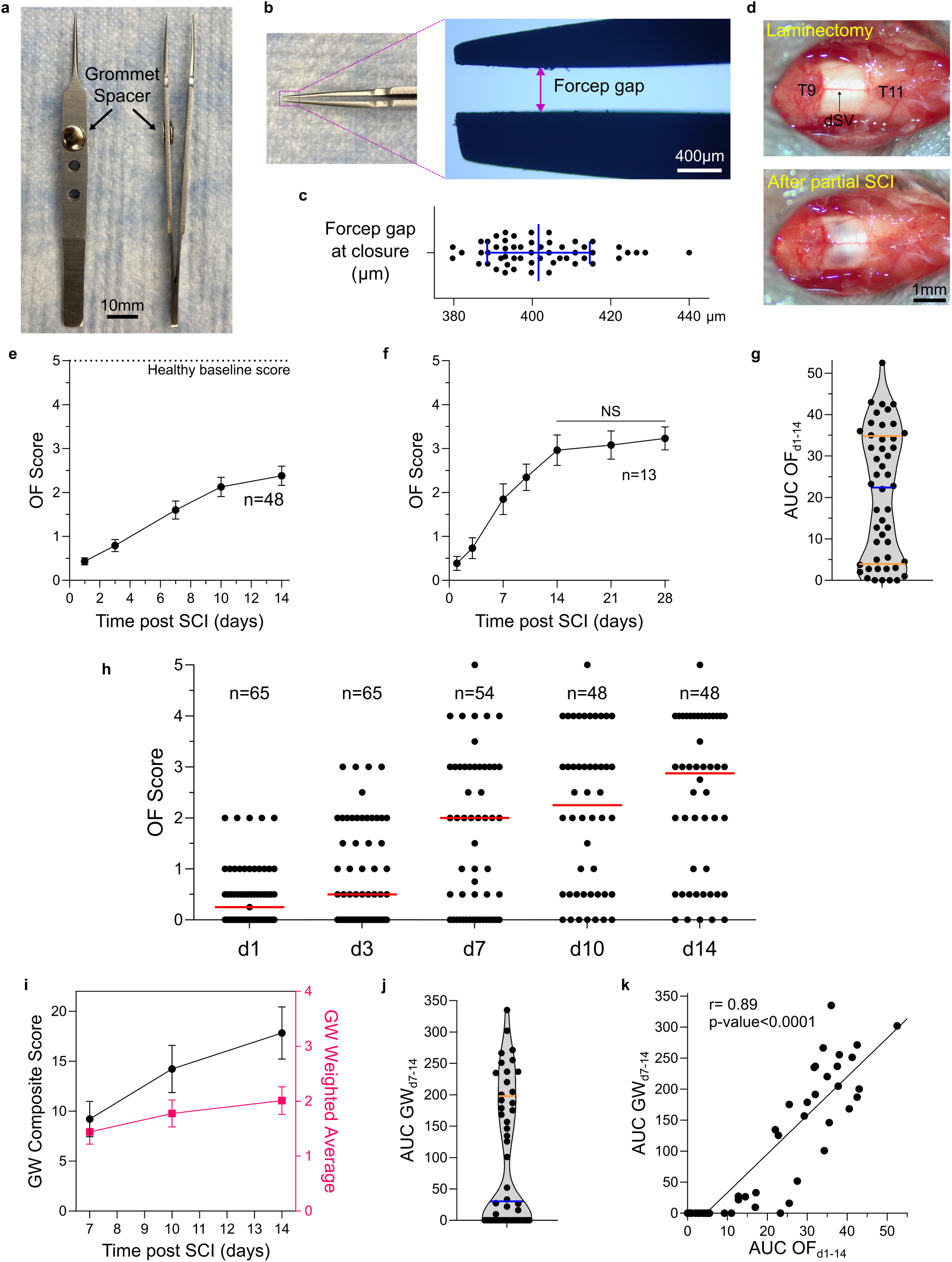
Population analysis of motor recovery in a partial crush SCI model. **a-c.** Overview and characterization of partial crush forcep tool including the average gap after a large number of forcep closure trials (**c**). **d.** Stereomicroscope images of a mouse spinal cord after laminectomy at T10 and application of the partial crush SCI (dSV = dorsal spinal vein). **e.** Mean Open field (OF) locomotion scores for the entire study population (n=48) over two weeks. **f.** Mean Open field (OF) locomotion scores across a cohort of animals evaluated over four weeks (n=13), No significant differences, NS, in OF score from 14-28 days, One-way ANOVA with Tukey’s multiple comparisons test. **g.** Area under the curve (AUC) of the OF score trace from days 1 to 14 (AUC OF_d1-14_) for the entire study population. **h.** Individual animal OF scores with population sample size indicated above each time point. Red bar represents the median of the population at the specific timepoint. **i.** Evaluation of grid walk (GW) test performance for the entire study population (n=48) at days 7, 10 and 14 post SCI. **j.** AUC of the GW composite score from days 7 to 14 (AUC GW_d7-14_) for the entire study population. **k.** Correlation analysis of OF and GW scores, with the behavioral tests showing strong positive correlation (r=0.89), p-value <0.001, t-test for Pearson’s linear correlation. Graphs in e, f, i show mean + standard error of mean (s.e.m.). Graph in c shows mean + standard deviation with individual data points overlayed. Graphs in g and j show median (blue) and quartiles (orange).

To evaluate locomotion recovery in the partial crush SCI we applied an open-field (OF) behavioral test using the simple 6-point scale that we and others have used extensively before, and which has demonstrated to be particularly useful for characterizing variations in wound repair outcomes at thoracic crush SCI caused by differences in injury severity^24^ or perturbations of astrocyte wound responses^3,25^. On this 6-point scale, a score of 5 equates to normal walking while a score of 0 is no movement of the hindlimbs of any kind^24^. The 6-point scale was used here because it is effective at assessing only gross voluntary hindlimb movement while also being more straightforward to implement and maintain consistency across different lab personnel when compared to other observer-scored scales, such as BBB or BMS, which require extensive specialized training and accrued experience to execute reliably^26^. We anticipated that recovery trajectories would be sufficiently assessable using the 6-point scale scoring system. For inclusion in the study, mice were required to show complete paralysis immediately after crush injury surgery, as well as not recovering any weight support capabilities on the first day after injury (i.e. open field day 1 score (OF_d1_) of ≤2). Analyzing the training data cohort of 48 mice that were evaluated over a 14 day period we noted, as expected^3^, a gradual and substantial improvement in locomotion recovery during the first 10 days post injury across the population with OF scores plateauing by 14 days (**Figure 1e**). No significant improvement in motor function beyond this 14-day plateau value was noted for a smaller cohort of 13 mice that were evaluated over an extended 28-day period (**Figure 1f**). Taking the area under the curve of the OF score trace from days 1 to 14 (AUC OF_d1-14_) generated a single relative recovery metric. Mice progressing to higher OF scores at earlier post injury times and to higher plateau levels achieved greater AUC OF_d1-14_ values (**Figure 1g**). By the AUC OF_d1-14_ metric and by inspection of individual animal scores on the temporal OF score trace (**Figure 1h**), there was a marked spread of behavioral outcomes across the population, with some mice recovering quickly while others not at all. The greatest spread in behavioral recovery was noted at the 7-day post SCI timepoint. There was a median OF score of 2.875 on post injury day 14 but there were nearly as many mice with a score of 1 or less (15/48) as there were mice obtaining a score of 2-3 (17/48), or 4+ (16/48) at this timepoint indicating substantially disparate functional capabilities across the population (**Figure 1h**).

To further characterize functional deficits in motor coordination, proprioception, fine locomotor control, and sensorimotor integration, we tested the same partial SCI crush mice on a grid walk (GW) apparatus. Since evaluation on the GW test requires a capacity for weight-supported stepping, we assessed animals from 7 days onwards when many of the cohort had started to recover such functions. To evaluate performance on GW, we derived a composite score for each testing session that integrated both the quality of stepping events (a weighted average) and the number of observed plantar placement stepping events, using a multiplicative formulation. We assigned successful plantar placed steps on grid rungs with a weighting of 4, slight slips off the grid with a weighting of 2, and total missed steps through the grid with a weighting of 1, and used these to compute the weighted average assessment of stepping quality during the 3-minute testing session. The weighted average metric rewarded mice that made fewer stepping mistakes during the testing session, while the composite score incorporated a multiplicative factor that was the square root of the total number of successful plantar placed steps, which elevated mice that were more active during the testing. Mice that could not perform any steps on the test received a score of 0. We observed a steady temporal increase in both the weighted average and composite score across the population from days 7 to 14 (**Figure 1i**). Taking the AUC for the composite score from day 7 to 14 could be used to establish a single outcome recovery metric for the GW test (AUC GW_d7-14_) (**Figure 1j**). Just like for the OF testing, there was considerable spread in AUC GW_d7-14_, but there was a strong positive correlation between AUC OF_d1-14_ and AUC GW_d7-14_ with a Pearson’s correlation (r) of 0.89, suggesting a general concurrence between the performance of the partial crush SCI mice on the two tests (**Figure 1k**).

These data show that mice subjected to a partial crush SCI using our newly constructed forcep tool show a natural recovery trajectory that involves a steady functional improvement that plateaus by 14 days post injury. Despite this, there is notable variation in the natural recovery trajectory across the population, with some mice recovering stepping functions quickly and some mice showing only limited hindlimb movement by 14 days, highlighting the need for analytical approaches to classify mice into more appropriate subgroups for analysis.

### Classifying partial SCI mice into subgroups using acute data enables open field recovery predictions

From our experiences with the partial crush model, we have noted 3 general categories of recovery profiles across animals that we are designating as Class 1, 2, and 3 recovery trajectories for the purposes of this work. **Class 1** animals show limited recovery and usually plateau to a recovery level of extensive hindlimb motion without the ability to engage in weight support or stepping. **Class 2** animals recover slowly over 14 days but reach plateauing recovery levels characterized by stepping capabilities with obvious and persistent disability compared to healthy baselines. **Class 3** animals recover very quickly (within a week or less) but reach similar recovery plateaus as Class 2 animals. We wanted to see if we could identify and classify mice displaying these 3 distinct classes of recovery using only data that could be collected during the post-injury period before experimental treatments would be administered. To do this, we used the OF score results taken on post injury days 1 and 3 to generate a single acute functional metric by taking the AUC between the two timepoints (AUC OF_d1-3_). To simplify nomenclature, we renamed this AUC OF_d1-3_ parameter as the Acute Functional Score (AFS) (**Figure 2a**). We avoided incorporating grid walk testing data or kinematic assessments to generate the AFS for our subgroup categorization since the information derived from these tests at acute post injury timepoints is limited, with mice generally lacking weight support or stepping capabilities. There was a strong positive correlation between AFS and both AUC OF_d1-14_ (r = 0.85) (**Figure 2b**) and AUC GW_d7-14_ (r = 0.82) (**Figure 2c**), suggesting that the early behavioral outcomes used to define the AFS are adequate predictors of the overall recovery in individual mice.

**Figure 2:**
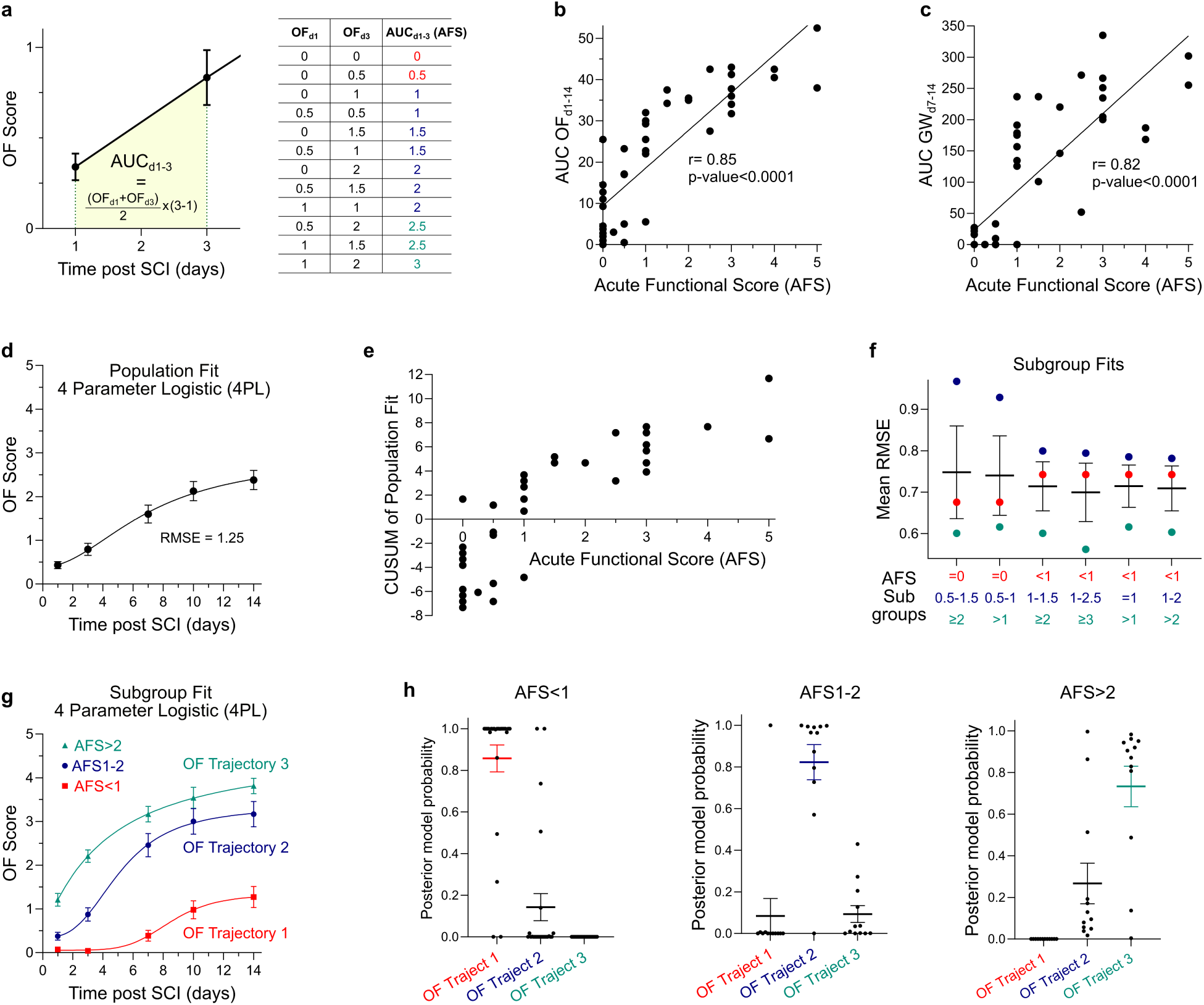
Classifying subgroups from open field (OF) behavioral testing to predict functional recovery after partial crush SCI. **a**. Overview of AFS calculations that involve computing the area under the curve (AUC) of the OF trace from post SCI days 1 to 3. **b.&c.** Correlation analysis of AFS with OF (**b**) and GW (**c**) scores, with the AFS values showing strong positive correlation with both OF (r=0.85) and GW (r=0.82) results, p-value <0.001, t-test for Pearson’s linear correlation. **d.** 4 parameter logistic (4PL) fit to population mean OF results. **e.** Cumulative sum residuals (CUSUM) for the 4PL population fit for each mouse related to its AFS designation. **f.** Mean root mean square error (RMSE) for six unique, non-overlapping arrangements of three AFS subgroupings. **g.** 4 parameter logistic (4PL) fitted to OF data for the subgroups AFS<1, AFS1-2, and AFS>2 to derive OF trajectories 1, 2, and 3. **h.** Posterior model probabilities (PMP) for the fits for each of the three OF trajectories for each mouse separated by AFS designation. Graphs in d,f, g, h show mean + standard error of mean (s.e.m.) with individual data points overlayed.

Although the OF test uses a scale of ordinal integers, we treated OF scores as quasi-continuous for curve fitting and trajectory modeling purposes, since the underlying functional recovery processes exist on a latent continuum that is monotonically progressing and the intervals of the OF test scale are approximately equally spaced^27^. By treating the data in this way, and given the noted trend of recovery, we fitted the population with a four-point parameter (4PL) sigmoidal curve as part of our initial latent growth curve analysis (**Figure 2d**). Using this fitted equation, we could readily estimate the mean plateau OF score (2.90) and the mean timepoint of recovery acceleration (inflection point) (7.04 days) for the population. The root mean square error (RMSE), a measure of the regression model’s predictive performance, for the population fit was 1.25, which was only moderately better than the population standard deviation across all timepoints of 1.46. Cumulative sum residuals (CUSUM) were calculated to evaluate the deviation from the fitted regression on a per animal basis (**Figure 2e**). Plotting CUSUM against AFS values for each animal showed that mice with low AFS were generally overestimated by the fitted regression (CUSUM<0), whereas mice with high AFS were substantially underestimated by the fitted regression (CUSUM>0) (**Figure 2e**). These results suggest that separating the population into sub-groups may improve the recovery trajectory fit. To identify the most appropriate AFS subgroup classifications for further analysis, we re-fit separate 4PL regressions to different sets of three non-overlapping AFS subgroups and used the mean and range of RSME values for the three regressions to quantitatively assess the suitability of the subgroupings (**Figure 2f**). Of the six unique, non-overlapping arrangements of three AFS subgroups, the subgroupings of AFS<1, AFS1-2, and AFS>2 showed one of the lowest RMSE means, smallest RMSE variance, and lowest maximum RMSE values among the subgrouping options, making it most appropriate for further investigation (**Figure 2f**). As expected, there were lower RMSE values for all subgroups when fitted to their uniquely defined trajectory regressions (mean RMSE = 0.71) than for the population fit (RMSE = 1.25), suggesting an overall improvement in goodness-of-fit with the subgroup designations (**Figure S1**).

For the selected AFS subgroup designations, out of a population of 48 mice, there were a total of 24 mice in AFS<1, 12 mice in AFS1-2, and 12 mice in AFS>2. The 3 distinct regressions showed good fit for the subgroup means (**Figure 2g, S1**). We designated the regressions for the 3 subgroups as OF Trajectories 1, 2 and 3 with the numbers chosen to match the 3 classes of recovery profiles (Class 1, 2, 3) we defined above since the recovery trajectories generally matched the noted phenotype of these recovery classes. Based on the parameters of the fitted 4PL regression, Trajectory 1 had a substantially lower plateau OF score (1.34) compared to Trajectory 2 and 3 (plateau OF score = 3.34 and 4.67 respectively), which is characteristic of Class 1 recovery whereby mice regain some hindlimb movement but not the ability to step. Trajectory 2 and 3 differed in their inflection point values (5.14 versus 4.20 days) as well as the time taken to reach the stepping OF score of 3 threshold (10.40 versus 5.98 days), reflecting the slower rate of recovery characteristic of Class 2 and the faster rate of recovery seen in Class 3. To assess the uniqueness of the three recovery trajectories and the AFS designations that define them, we determined how well each animal’s individual recovery profile was described by any one of the three trajectories using leave-one-out cross-validation (LOOCV) and computing posterior model probabilities (PMP). This analysis involved calculating for each mouse the residuals for the fits for the three trajectories using fitted regressions derived from a version of the dataset that did not include the specific animal being analyzed (**Figure 2h**). Or put another way, when assessing an individual mouse against the three regressions, we did not use that mouse’s OF data to derive the regressions. Thus, the PMP values reflect the relative evidence for the suitability of any one of the three fitted trajectory regressions to describe an individual animal’s OF recovery. Mice with an AFS<1 had a mean PMP of 0.86, 0.14, and 0 for OF Trajectory 1, 2 and 3 respectively, suggesting that Trajectory 1 was at least 6 times more probable for AFS<1 mice on average than any of the other two trajectories. AFS1-2 mice had a high mean PMP (0.82) for OF Trajectory 2, which made it over 8 times more probable than either of the other two trajectories, which both had PMPs less than 0.1. AFS>2 mice were described by OF Trajectory 3 at a mean PMP of 0.73, providing moderately strong confidence that these mice will undergo Class 3 recovery and not Class 2 recovery which had a mean PMP of 0.27 for AFS>2 animals. Overall, 40 of the 48 mice (83.3% of the cohort) could be suitably estimated by the OF trajectory that was associated with their AFS designation (**Figure 2h**). However, while the trajectories generally described the AFS defined subgroups adequately, 4/24 animals with AFS<1 had a higher probability of being better described by OF Trajectory 2, 1/12 animals with AFS1-2 were better assigned to OF Trajectory 1, and 3/12 animals with AFS>2 were better suited to OF Trajectory 2, suggesting that a small number of mice either had greater deficits than the AFS revealed or they recovered more quickly than most of the other mice with similar AFS designations. Nevertheless, it was notable that there were no mice in the AFS>2 subgroup that were described best by OF Trajectory 1, and no mice in AFS<1 had probabilities greater than 0.002 for OF Trajectory 3, suggesting an essentially certain distinction between these two AFS defined subgroups by the OF evaluations.

These data show that OF data captured during the first 3 days post partial SCI can be used to estimate the recovery trajectory on this behavioral test over 14 days for individual mice with a strong level of confidence. More importantly, the AFS definitions can be used to distinguish mice that will rapidly recover versus those that will undergo very little spontaneous recovery with essentially 100% accuracy.

### Incorporating grid walk assessments improves the understanding of recovery trajectories

Since the AFS subgroup classification effectively differentiated the three classes of recovery on OF testing and given the strong overall correlation between OF and GW tests across the population dataset, we next evaluated whether the acutely defined subgroups could also capture differences in spontaneous recovery outcomes on the GW test (**Figure 1j**). Separating the mice into the same AFS subgroup designations as defined by the OF evaluations revealed significant differences in GW outcomes at 7, 10 and 14 days, as assessed by total numbers of plantar placement stepping events (**Figure S2**), GW weighted average (**Figure 3a**) and GW composite score (**Figure 3b**), consistent with the recovery class. Specifically, AFS>2 mice exhibited early recovery with high-scoring GW results at the 7-day evaluation relative to the other subgroups; AFS1-2 mice showed delayed recovery that ultimately matched the faster recovering AFS>2 animals by 14 days; and AFS<1 mice demonstrated essentially no recovery on the GW test. These temporally dependent differences on the GW test produced clear separation by the single GW outcome recovery metric, AUC GW_d7-14_, between all three AFS subgroups (**Figure 3c, S2**).

**Figure 3:**
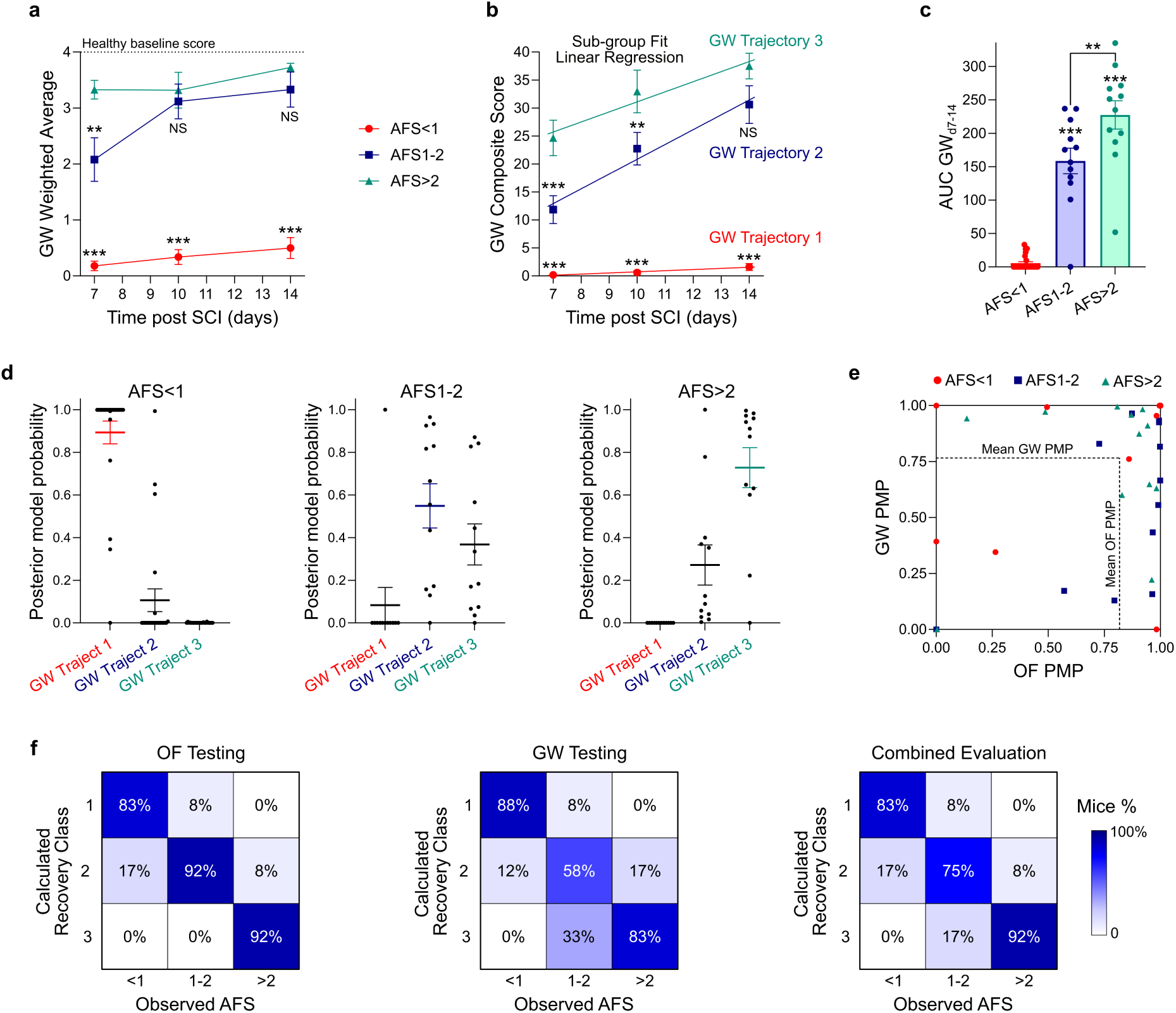
Incorporating gridwalk (GW) assessments into post SCI recovery predictions. **a**.**&b.** GW weighted average (**a**) and GW composite scores (**b**) for the subgroups AFS<1, AFS1-2, and AFS>2. *** p-value<0.0001, ** p-value<0.004, Not significant (NS) versus AFS>2 subgroup at the corresponding timepoint, Two-way ANOVA with Tukey’s multiple comparisons test. GW composite scores across the three subgroups were fitted with simple linear regressions to derive GW Trajectories 1, 2, and 3. **c.** AUC of the GW composite score from days 7 to 14 (AUC GW_d7-14_) for the AFS subgroups. *** p-value <0.0001, ** p-value<0.004, Not significant (NS), One-way ANOVA with Tukey’s multiple comparisons test. **d.** Posterior model probabilities (PMP) for the fits for each of the three GW trajectories for each mouse separated by AFS designation using leave-one-out cross-validation (LOOCV). **e.** Comparison of OF PMP and GW PMP values for the AFS defined trajectory for each mouse in the population (n=48). **f.** Confusion matrices showing the percentage of mice displaying Class 1, 2 and 3 type recovery from the three AFS subgroups based on OF testing, GW testing, and the combined evaluation of the two tests. Graphs in a-d show mean + standard error of mean (s.e.m.) with individual data points overlayed.

Based on the trend of GW composite scores temporally across the three subgroups, we fitted these data with a simple linear regression to generate GW Trajectories 1, 2 and 3 (**Figure 3b**), before computing GW PMPs for each AFS subgroup using LOOCV, consistent with the modeling approach described in the prior section for the OF data analysis (**Figure 3d**). Mice with an AFS<1 showed a high mean PMP for GW Trajectory 1 (0.89) and correspondingly low support for both GW Trajectories 2 (PMP=0.11) and 3 (PMP=0.001). For AFS1-2 mice, the associated GW Trajectory 2 had a moderate mean PMP of 0.55, only just outperforming GW Trajectory 3 (PMP=0.37). For AFS>2 mice, the GW Trajectory 3 was more strongly supportive (PMP=0.73), with a lower probability for GW Trajectory 2 (PMP=0.27). As in the OF trajectory evaluations, no AFS>2 mice were better described by GW Trajectory 1, nor were any AFS<1 mice better fit by GW Trajectory 3. Although on average both AFS1-2 and AFS>2 subgroup mice showed a preference for their own AFS-defined trajectory, there was substantial cross-group support for both trajectories. This lower confidence is likely because there are fewer sampled timepoints for the GW test compared to the OF evaluation, since GW data collection started at 7 days rather than 1 day after injury, as was the case for the OF data. Furthermore, the two fitted trajectory regressions converge by 14 days meaning that practically only day 7 and day 10 GW results distinguished the two subgroups. Nevertheless, these results show that overall, the AFS designations do permit effective recovery class distinction for the GW test with strong confidence.

Because OF and GW test results were strongly correlated at the population level, it was unsurprising that most mice (34/48 or 70.8%) had PMP values greater than 0.5 for the AFS-associated trajectory for both OF and GW (**Figure 3e**). However, a subset of mice (10/48) showed divergent results across the two tests, with a PMP greater than 0.5 for one test but not the other, which we attributed to the underlying complexity and inherent variability in spontaneous animal behavior and its observation (**Figure 3e**). To increase robustness, and because the two metrics are strongly correlated and therefore not independent, we computed a combined evaluation metric by taking the average of the OF and GW PMPs (referred to henceforth as the combined PMP). Each mouse was then assigned a recovery class according to its combined PMP for the associated trajectory and summarized in a confusion matrix (**Figure 3f**). The combined PMP metric provides a conservative and practical approach for integrating information from the two behavioral tests, setting a deliberately high threshold for identifying deviations from the natural post SCI recovery trajectory, requiring substantial differences in recovery outcomes on both tests for the recovery class to be re-assigned. Such an approach is intended to minimize the potential for exaggerating the efficacy of tested experimental interventions. The combined evaluation showed that AFS designations strongly predicted the eventual recovery class for most animals, with 83%, 75% and 92% correct assignments for AFS<1, AFS1-2, and AFS>2 mice, respectively (**Figure 3f**). Flattening the confusion matrix into a 9-element vector and calculating its cosine similarity (CS) to an ideal identity matrix yielded a value of 0.98, which indicated strong correspondence (a CS cut-off of >0.9 was used as evidence of strong class-wide agreement). Notably, the combined evaluation matched the predictive strength of OF testing alone (CS = 0.99) and improved on GW testing alone (CS = 0.94). Using the combined approach, only a small number of mice shifted to higher (6/48) or lower (2/48) recovery classes than predicted by the AFS designation (**Figure 3f, S3**).

These data show that AFS designations predict natural functional recovery outcomes on GW test after partial SCI with improved confidence for AFS<1 mice but moderately reduced confidence for AFS1-2 or AFS>2 mice when compared to predictions generated solely using the OF testing data. Nevertheless, incorporating results from both OF and GW tests improves the robustness of the recovery trajectory prediction for all three AFS-defined subgroups.

### Kinematics reveals locomotion differences across the three partial SCI recovery classes

To test whether differences in locomotion recovery between the three recovery classes could also be detected using more sensitive evaluations of gait and hindlimb coordination, we conducted markerless hindlimb kinematics analysis from a sagittal viewpoint on a small cohort of partial crush mice on a mouse-sized treadmill on days 7 and 14 post injury. Healthy and SCI mice that recovered hindlimb stepping capacity demonstrated effective walking on the treadmill when run at a speed of 7cm/s, with this speed maintained for all tested animals. We employed DeepLabCut and the Automated Limb Motion Analysis (ALMA) deep-learning toolbox^28^ to track hindlimb movements and measure important locomotion parameters during the gait cycle (**Figure 4a**). Stratification of partial crush SCI animals by their class of recovery, as defined by the individual animals’ combined PMP derived by integrating both OF and grid walk assessments, revealed notable qualitative differences in the overall kinematic profiles by video recording and the associated stick model plots that reflected the nature of the recovery class described above (**Figure 4b-d**). Class 1 mice showed pronounced hindlimb dragging at 7 days followed by some moderate improvements in recovery by 14 days, but with hindlimb dragging persisting. By contrast, Class 2 mice showed less severe hindlimb dragging at 7 days compared to Class 1 mice and displayed recovery of weight-supported stepping by 14 days. Class 3 mice rapidly recovered weight-supported stepping by 7 days that persisted through 14 days. In both Class 2 and Class 3 mice, their weight-supported stepping profiles at 14 days still showed some notable differences from those of healthy uninjured mice, particularly in the swing phase of the gait cycle (**Figure 4c,d**). To quantitatively evaluate the differences in locomotion between healthy and SCI mice, we assessed the variation in 31 parameters measured across the gait cycle by dimensionality reduction using principal component analysis (PCA) (**Figure 4e**). The two principal components from this analysis accounted for approximately 50% of the total variation for a system containing 23 individual kinematic assessments (6x healthy, 7x 7d SCI, and 10x 14d SCI mice) and revealed obvious temporal and recovery class-dependent effects on locomotion function (**Figure 4f,g**). PC1 defined severe locomotion dysfunction consistent with loss of stepping capacity, which was reflected by high factor loading contributions from parameters assessing movement of ankle and metatarsophalangeal (MTP) joints, time in stance phase, and the extent of hindlimb dragging (**Figure 4h, S4**). Notably, PC1 values correlated with AUC OF_d1-14_ scores (r=-0.68) but not AUC GW_d7-14_ scores, suggesting a general concurrence for the kinematics with another locomotion assessment (OF testing) but not with evaluations of sensorimotor integration (GW testing) (**Figure S4**). Since Class 1 mice failed to recover stepping functions, they showed significantly larger PC1 values at both 7 and 14 days post injury compared to both healthy controls and to other less severely injured SCI mice from the other subgroups (**Figure 4f**). At 7 days, Class 2 mice showed PC1 values that were significantly larger than both healthy and Class 3 mice. Recovery of stepping capacity in Class 2 mice by 14 days resulted in a reduction in PC1 values such that they were no different from Class 3 or healthy mice. The PC1 values for Class 3 mice were not significantly different from those of healthy controls at either days 7 or 14, consistent with the rapid return of stepping functions in these animals. PC2 defined differences in quality of recovered stepping functions in SCI mice compared to healthy controls with parameters for overall leg (toe-to-illiac crest) and knee positioning during gait as well as gait cycle speed having high factor loading contributions (**Figure 4h, S4**). Notably, at 14 days post injury, Class 2 and Class 3 mice had PC2 values that were indistinguishable from each other, but significantly higher than those of healthy controls (**Figure 4g**). These PC2 results corroborated the findings from the other behavioral tests that showed that Class 2 and Class 3 mice are indistinguishable from each other at 14 days but that both have persistent and detectable locomotion deficits compared to healthy uninjured mice.

**Figure 4:**
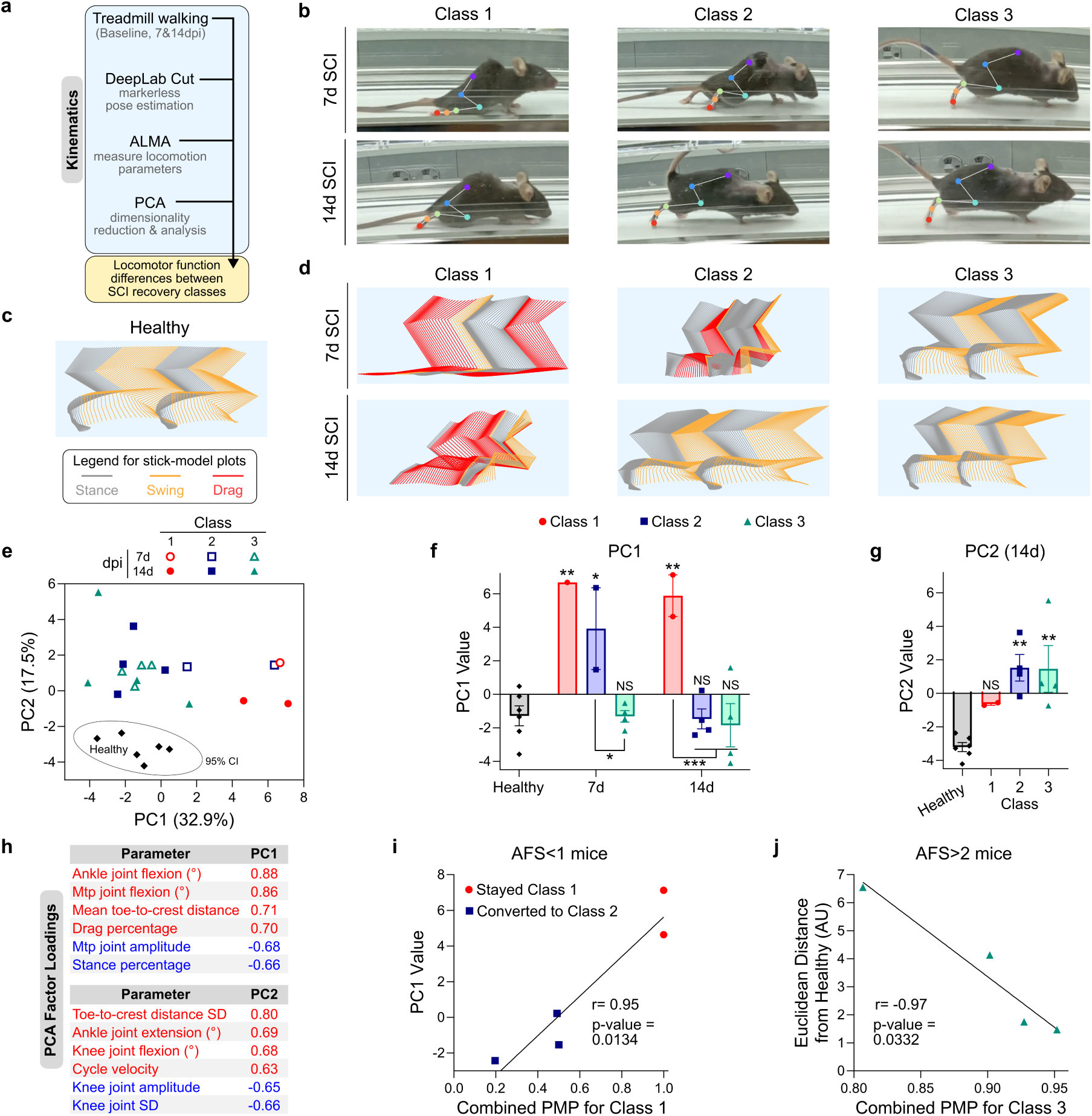
Treadmill based hindlimb kinematics assessments of mice from the three partial SCI recovery classes. **a.** Outline of kinematic analysis workflow. **b.** Images of representative mice from the three recovery class designations walking on the treadmill at 7 and 14 days with labeling of hindlimb joints. **c&d.** Stick-model plots for healthy (c.) and SCI mice (d.) during two gait cycles. **e.** Principal component analysis of 31 kinematic parameters for uninjured healthy and pSCI mice at 7 and 14 days. The ellipse represents the 95% CI for the healthy group. **f.** PC1 values for healthy and pSCI mice separated by post injury day and recovery class. ***p-value <0.001, ** p-value <0.01, * p-value<0.05, Not significant (NS), Two-way ANOVA with Tukey’s multiple comparisons test. **g.** PC2 values for healthy and pSCI mice at 14 days post injury separated by recovery class. ** p-value <0.01, Not significant (NS), One-way ANOVA with Tukey’s multiple comparisons test. **h.** Factor loadings defining PC1 and PC2. **i.** Correlation analysis of PMP and PC1 values for AFS<1 mice, with PMP values showing strong positive correlation (r=0.95) with the PC1 values allowing for the distinction of class converting and persisting AFS<1mice, t-test for Pearson’s linear correlation. **j.** Correlation analysis of Euclidean distance from healthy and Class 3 PMP values for all AFS>2 mice showing strong negative correlation (r=-0.97) between the Euclidean distance and the PMP value, t-test for Pearson’s linear correlation. Graphs in f&g show mean + standard error of mean (s.e.m.) with individual data points overlayed.

In the cohort of mice undergoing treadmill kinematics assessments, we evaluated 5 mice designated as AFS<1 based on their acute OF evaluations. However, three of these AFS<1 mice were deemed to have converted to Class 2 recovery because their combined PMP was less than 0.5 for the Class 1 recovery trajectory and highest for the Class 2 recovery trajectory, meaning they recovered better than predicted by the initial AFS designation. Thus, we used kinematics data from these three Class 1 to 2 converting mice to determine whether the potentially more sensitive kinematics results correlated with the combined OF and GW trajectory prediction. Plotting the PC1 values from the kinematics against the combined PMPs for Class 1 recovery for all AFS<1 mice including both the Class 1 to 2 converting and Class 1 persisting animals showed that there was strong positive correlation (r=0.95) between the kinematics and combined OF/GW trajectory assessment (**Figure 4g**). This analysis showed that Class 1 to 2 converting mice had both lower PC1 and combined PMP values whereas Class 1 persisting mice, that maintained greater locomotor dysfunction, showed the opposite trend. In a similar manner, comparing the Euclidean distance from healthy mice on the PCA for all Class 3 mice (all AFS>2) against their combined PMP for Class 3 recovery revealed that mice that had the strongest predicted probability for Class 3 recovery (highest PMP) had the most similar locomotor capabilities to healthy mice on the kinematics (smallest Euclidean distance). Together these correlation assessments indicate that there was a strong correspondence among the three different motor behavioral evaluations, while also demonstrating the robustness of the subgroup classification methodology.

Overall, these data show that treadmill based hindlimb kinematics assessments are effective at distinguishing functional locomotor differences between the 3 recovery classes of partial crush SCI mice and that kinematics assessments in conjunction with the OF and GW tests can be used to determine whether an individual animal deviates from its AFS-defined recovery trajectory.

### Subgroup classification distinguishes conferred injury severity and extent of astrocyte bridging at partial SCI lesions

Prior work employing the partial crush SCI model has shown that functional recovery outcomes assessed by behavioral testing correlate strongly with injury severity as assessed by histology^1,2,4,25^. In particular, the persistence of larger non-neural lesion cores filled with peripherally derived immune and stromal cells confers worse functional recovery after SCI, whereas lesions displaying greater volumes of spared neural tissue or wound repair astrocyte bridging have enhanced recovery outcomes^1,2,4,25,29^. To evaluate whether the AFS subgroup classifications revealed significant differences in these key lesion severity metrics, we performed immunohistochemistry on horizontal partial crush SCI tissue sections taken at 3 days (**Figure S5**) and 14 days (**Figure 5a**) post injury, quantifying Gfap-positive astrocyte bridging (**Figure 5b-d**) as well as Cd13-positive, Gfap-negative non-neural lesions (**Figure 5e-g**).

**Figure 5:**
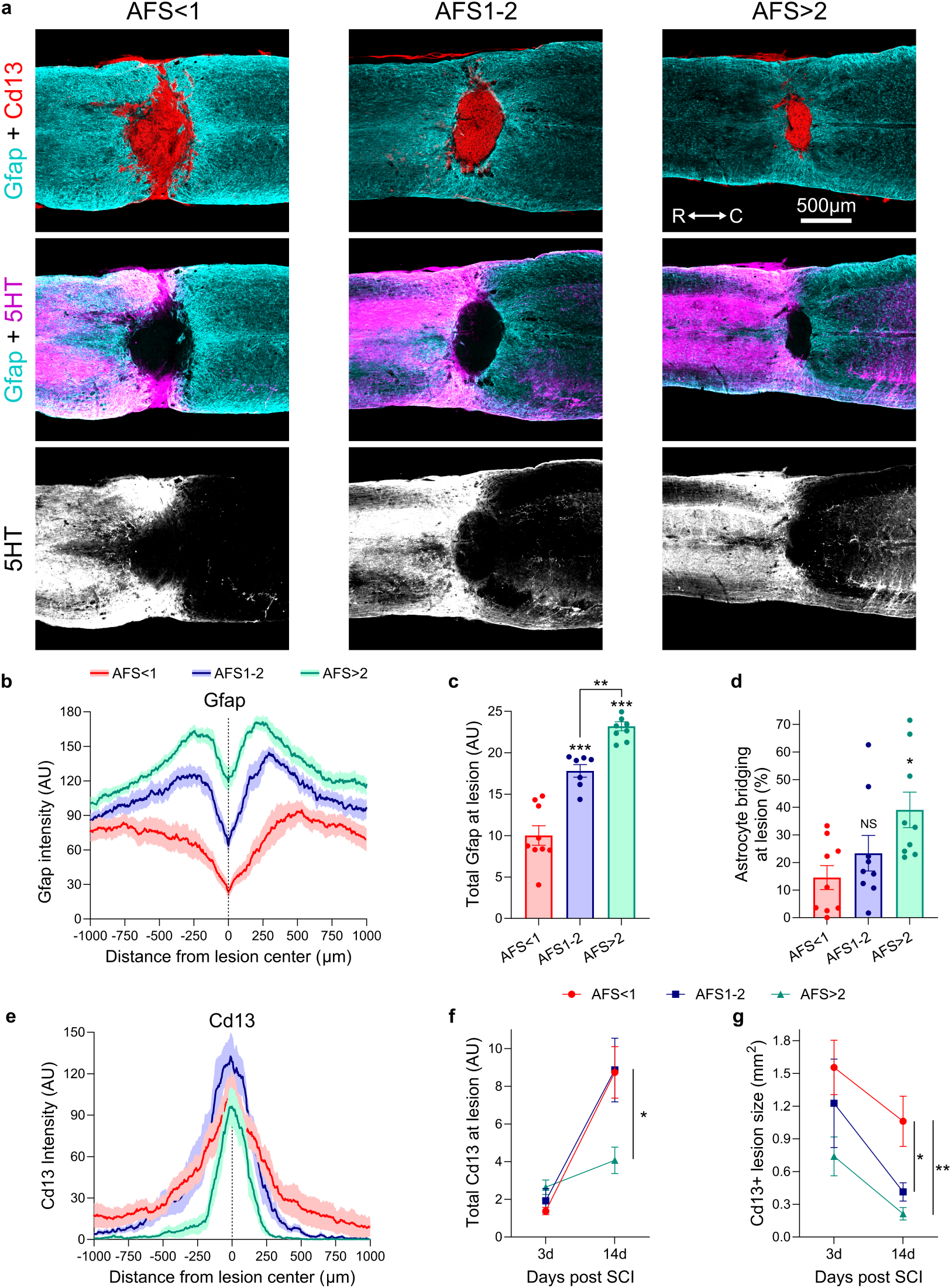
Subgroup classification distinguishes conferred injury severity and extent of astrocyte bridging at partial SCI lesions. **a.** Representative survey IHC images of SCI lesions from the three AFS defined subgroups at 14 days after partial crush injury showing Gfap-positive astrocytes, Cd13-positive immune and stromal cells within the lesion core, and 5HT-positive descending serotonergic tract fibers. **b.** Gfap intensity plots for partial SCI mice stratified by subgroup. **c.** Quantification of total Gfap via an area under the curve (AUC) calculation of Gfap intensity from 500µm rostral to 500µm caudal of the lesion epicenter showing a subgroup dependent increase in total Gfap. ***p value < 0.0001, **p value < 0.002, One-way ANOVA with Tukey multiple comparison test. **d.** Quantification of extent of astrocyte bridging at SCI lesions stratified by subgroup, *p value < 0.02, One-way ANOVA with Tukey multiple comparison test. **e.** Cd13 intensity plots for partial SCI mice stratified by subgroup. **f.** Quantification of total Cd13 at SCI lesions at 3 and 14 days after SCI stratified by subgroup. *p value < 0.05, One-way ANOVA with Tukey multiple comparison test on 14d data **g.** Quantification of SCI lesion size at 3 and 14 days after SCI stratified by subgroup. **p value < 0.002, *p value < 0.05, One-way ANOVA with Tukey multiple comparison test on 14d data. Gfap and Cd13 intensity plots show mean as darken lines and s.e.m as shaded area. All graphs are mean ± s.e.m.

At 14 days, SCI lesions from all subgroups contained high cell density Gfap-positive astrocyte borders derived from newly proliferated astrocytes^30^ that organized around Gfap-negative, Cd13-positive non-neural lesion cores located medially within the affected spinal cord tissue segment (**Figure 5a,b**). The total number of Gfap-positive astrocytes within a fixed 1mm width segment centered on the lesion varied significantly across the three subgroups of mice and scaled proportionally with recovery class (**Figure 5c**). Poorly recovering AFS<1 mice had the lowest number of Gfap-positive astrocytes, while rapidly recovering AFS>2 mice had over two-fold more Gfap-positive astrocytes within the SCI lesion segment (**Figure 5c**). AFS<1 mice had the largest sized Gfap-negative, Cd13-postive lesions compared to the other two subgroups at both 3 days and 14 days SCI (**Figure 5e,g, S5**), with astrocyte bridging covering less than 15% of the spinal cord cross section on average at 14 days and some AFS<1 mice having no detectable astrocyte bridging at all (**Figure 5d**). Likely due to this limited astrocyte bridging, there was a complete loss of descending serotonergic (5HT) tract fibers below the level of the injury and negligible spontaneous regeneration of these axons across the lesion in AFS<1 mice (**Figure 5a**). AFS1-2 mice had non-neural lesion cores (0.41mm^2^) that were less than half the cross-sectional area of AFS<1 mice (1.06 mm^2^) at 14 days (**Figure 5g**), while the astrocyte bridging accounted for nearly 25% of the spinal cord cross section in AFS1-2 mice. This enhanced astrocyte bridging correlated with increased 5HT axon sprouting in spinal cord tissue caudal to the lesion (**Figure 5a**). Interestingly, the AFS1-2 mice, despite having dramatically smaller lesions than AFS<1 mice, showed essentially equivalent total Cd13-positive cell recruitment into lesion cores by 14 days post SCI (**Figure 5f**), reflecting an increased density of these peripherally derived cells within a more organized, volume-constrained lesion. As anticipated, AFS>2 mice had the smallest lesions (0.21mm^2^) of the three sub-group classifications, which was almost half the cross-sectional area of the AFS1-2 subgroup and nearly one-fifth the size of AFS<1 lesions. The differences in lesion severity across the three subgroups of mice were conferred acutely, with lesions at 3 days showing a similar AFS-defined size trend (**Figure 5g, S5**). The smaller lesions in AFS>2 mice resulted in reduced total recruitment of Cd13-positive immune and stromal cells (**Figure 5f**), as well as a near two-fold increase in the persistence of astrocyte bridges compared to AFS1-2 mice (**Figure 5d**). Likely because of the astrocyte bridging occupying nearly 40% of the spinal cord cross-section, AFS>2 mice had the greatest extent of 5HT fiber sprouting into the caudal spinal cord (**Figure 5a**). Intact myelin basic protein (Mbp)-positive tissue at the lateral margins of the tissue surrounding the lesion cores in AFS1-2 and AFS>2 mice suggested that the glial cell bridging was derived from spared surviving neural tissue rather than exclusively from enhanced mobilization of wound repair astrocytes (**Figure S5**).

These data show that the AFS subgroup classification distinguishes differences in the severity of partial crush SCI lesions by immunohistochemistry, with AFS<1 mice having the largest lesions and AFS>2 mice having the smallest lesions. AFS>2 mice showed robust astrocyte bridging across nearly half the width of the spinal cord at the lesion site, which resulted in enhanced persistence of descending axons caudal to the lesion, an anatomical feature likely necessary for the motor recovery observed in these mice. Notably, AFS<1 mice show limited astrocyte bridging at lesions and descending tract axons caudal to the lesion, highlighting the essential role of these cellular elements in effective functional recovery after SCI.

### Subgroup classification predicts natural recovery of vehicle treated SCI mice and can be used to identify procedural bias

As part of pilot studies in our group, we have generated several animals that have received vehicle treatments of saline or an unloaded coacervate biomaterial carrier injected into the lesion epicenter at 3 days post partial SCI. The coacervate biomaterial has been previously characterized and demonstrates excellent *in vivo* biocompatibility in the CNS^31^. We have no basis to suspect that these vehicle-treated mice would have recovery profiles different from those in the training data assessed above. However, based on initial analysis of the entire population of mice in the two vehicle groups, it emerged that coacervate-treated mice had pronounced recovery improvements on both OF and GW testing compared to the saline control (**Figure 6a,b**). As before, the OF and GW tests were strongly positively correlated (r=0.93), and the coacervate-treated mice had the four best recovery outcomes on both tests (**Figure 6c**).

**Figure 6:**
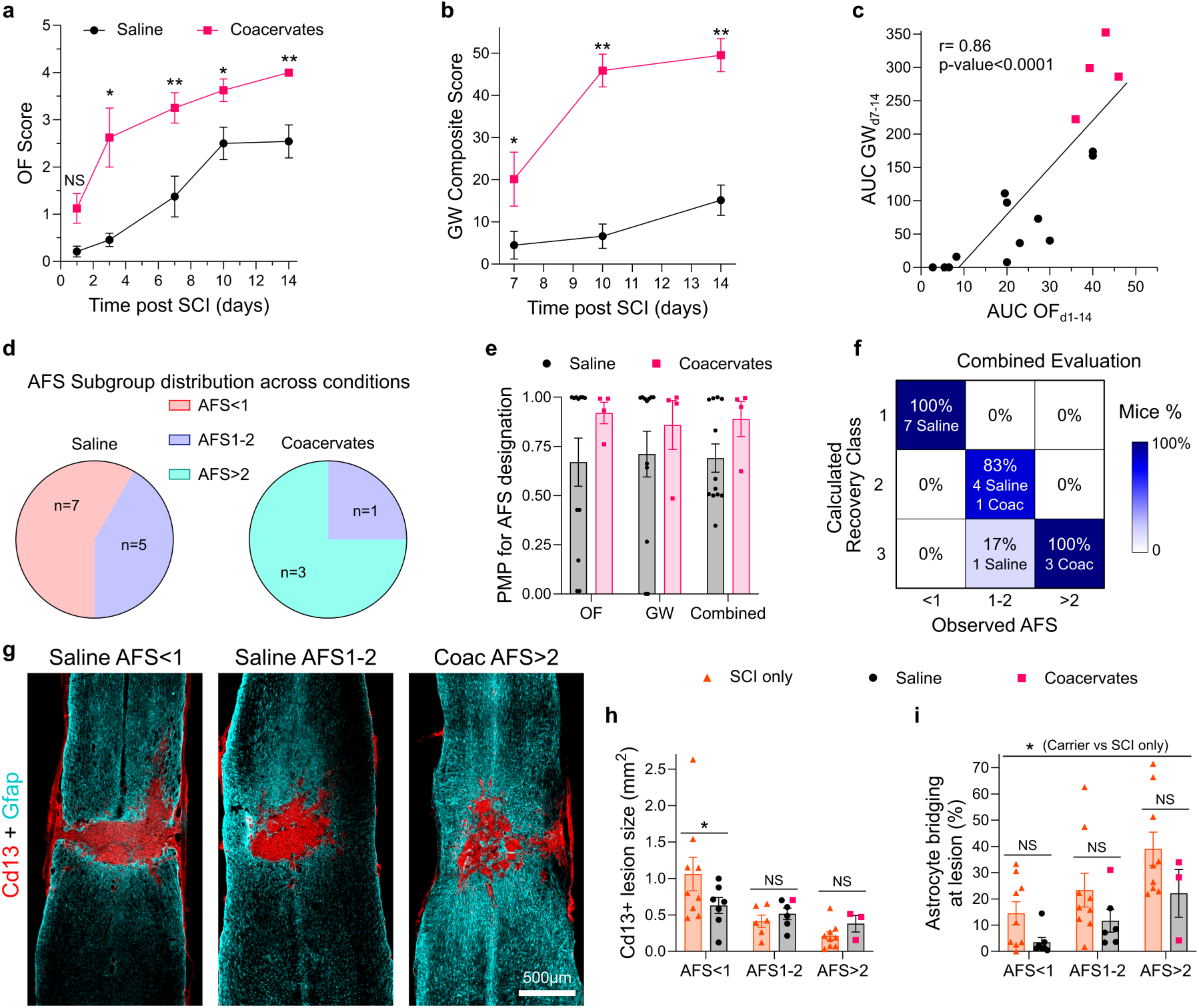
Subgroup classification predicts natural recovery of vehicle treated SCI mice. **a.** Mean Open field (OF) locomotion scores for the saline and coacervate treated SCI mice over two weeks. **p value < 0.005, *p value < 0.05, One-way ANOVA with Tukey multiple comparison test. **b.** Grid walk (GW) test performance for the saline and coacervate treated SCI mice over two weeks, **p value < 0.0001, *p value < 0.02, One-way ANOVA with Tukey multiple comparison test. **c.** Correlation analysis of OF and GW scores for carrier injected SCI mice, with the behavioral tests showing strong positive correlation (r=0.86), p-value <0.0001, t-test for Pearson’s linear correlation. **d.** Pie charts indicating the AFS subgroup distribution across the saline and coacervate injected SCI mice. **e.** Mean OF, GW and combined assessment PMP values for the AFS designation separated by carrier injection type. **f.** Confusion matrix showing the percentage of mice displaying Class 1, 2 and 3 type recovery from the three AFS subgroups based on the combined evaluation of OF and GW testing results. **g.** Representative survey IHC images of carrier treated SCI lesions from the three AFS defined subgroups at 14 days after partial crush injury showing Gfap-positive astrocytes and Cd13-positive immune and stromal cells. h. Quantification of SCI lesion size at 14 days after SCI stratified by subgroup and carrier injection. *p value < 0.05, Two-way ANOVA with Tukey multiple comparison test. **i.** Quantification of extent of astrocyte bridging at SCI lesions stratified by subgroup and carrier injection., *p value < 0.02, Two-way ANOVA with Tukey multiple comparison test.

To determine whether these significantly different recovery outcomes were due to a therapeutic effect from the unloaded coacervate carriers or to unintended procedural bias, we separated the two conditions into the three subgroups using the established AFS designations. We noted that within the saline-treated group, there were 7 and 5 mice belonging to the AFS<1 and AFS1-2 subgroups respectively. By contrast, the coacervate group had 3 mice in the AFS>2 subgroup and one mouse in the AFS1-2 subgroup, clearly showing that the coacervate group had a disproportionately higher representation of mice predicted to recover more quickly and to a great extent naturally without treatment application (**Figure 6d**). Computing OF and GW PMP, as well as the combined PMP derived from the average of the two tests, revealed that 11 of 12 mice in the saline group and 4 of 4 mice in the coacervate group (15/16 total or 93.8%) had recovery profiles that were suitably predicted by their AFS designation (**Figure 6e**). Only one mouse in the saline group deviated from its AFS-defined recovery trajectory, with PMP assessments suggesting it converted from Class 2 to Class 3 recovery, indicating its recovery trajectory was better than predicted by the AFS designation (**Figure 6f**). Ultimately, flattening the confusion matrix for the vehicle-treated mice and comparing it with an ideal identity matrix yielded a cosine similarity (CS) of 0.99, suggesting that vehicle-only injections did not meaningfully alter the recovery from the predicted natural trajectory (based on the same CS cut-off of 0.9 that we employed before). Evaluation of histological sections of SCI tissue from the saline and coacervate-treated mice corroborated the AFS designations (**Figure 6g**). The lesion sizes for the mice from the three AFS subgroups following carrier only injections were not significantly altered compared to untreated SCI mice for the AFS1-2 and AFS>2 subgroups, while there was a modest but significant reduction in lesion size for AFS<1 saline injected mice compared to the untreated benchmark (**Figure 6h**). Interestingly, the extent of astroglial bridging across the population of carrier-injected mice was overall lower than that of the untreated control, suggesting that the surgical injection alone at the 3-day post injury timepoint, in the absence of a loaded therapeutic agent, could interfere with natural astrocyte wound repair processes. However, Tukey’s post-hoc comparisons did not identify any specific subgroup in which astrocyte bridging was significantly altered, likely due to the somewhat high variance in this parameter across animals and the lower sample size for vehicle-injected mice.

These data show that AFS subgroup classifications made prior to administering experimental treatments can effectively identify procedural bias by revealing unequal injury severity distributions across experimental groups that would otherwise confound therapeutic recovery outcomes.

## Discussion

The partial crush spinal cord injury (SCI) paradigm is a useful preclinical model for studying CNS wound healing biology and evaluating the efficacy of new therapeutic strategies in mice, as neuroanatomical and behavioral recovery readouts can be either enhanced or diminished relative to controls through application of treatment or transgenic manipulation^2–6^. However, the positive attributes of the partial SCI model are also coupled with unavoidable, and often substantial, inter-animal variability in injury severity that is difficult to control or detect during experimentation, creating conditions for inadvertent procedural bias or high variance in behavioral recovery outcomes that can obscure or distort the true effects of experimental interventions^9,10^. To address this limitation, we developed a new analytical framework to predict the trajectory of natural, spontaneous post-injury functional recovery and overall lesion severity of partial SCI mice by incorporating sparse behavioral data collected acutely after SCI to define distinct recovery subgroups. Our findings demonstrate proof-of-principle that a subgroup classification approach is effective at predicting at least 3 distinct classes of behavioral recovery that occur naturally following partial SCI, while also clearly stratifying injury severity and extent of astrocyte bridging at partial SCI lesions as assessed by post-hoc histology. These findings have broad implications for the design of preclinical SCI studies to assess the efficacy of experimental treatments and establish a foundation for more advanced modeling approaches to further improve the prediction of natural post-SCI recovery trajectories in the preclinical setting.

Differences in injury severity following partial crush SCI can result in lesions with dramatically different histopathologic features, including the extent of hemorrhage, peripherally derived inflammation, demyelination, astrocyte bridging, descending axon tracts caudal to the lesion, among many other injury-associated features^32^. The current work showed that the population of partial SCI mice could be segregated into 3 subgroups that differed in astrocyte bridging, peripheral immune cell recruitment and density at lesion cores, as well as in the persistence of descending axons below the lesion. Experimental treatments which therapeutically target only a select few of these many SCI histopathological features, may show efficacy in certain lesion settings but fail in others^33^. Given the tremendous heterogeneity in SCI lesion location and severity within the human SCI population, establishing the robustness of a new experimental therapy across variable injury severity is essential. Although the inherent variability of the partial crush SCI model could be advantageous for assessing treatment robustness, no preclinical studies employing this model have been able to effectively assess the lesion severity prior to treatment on a per animal basis. As a result, studies are either severely underpowered due to high outcome variability, or worse, outcomes may be confounded by procedural bias because comparisons are made between control and treatment groups with different distributions of lesion severity^10^. Addressing these issues requires highly powered studies with large sample sizes, but this is becoming increasingly impractical in most pre-clinical research settings, with resources being more constrained than ever, and the ethical use and minimization of animals is mandated. The analytic framework developed here, which demonstrated an excellent capacity to distinguish lesions of differing injury severity using non-terminal data collected over the first 3 days post injury, will be useful in improving preclinical study design by minimizing intragroup variability, identifying and counteracting the impact of procedural bias (as seen in comparison between saline and coacervate vehicle treatments studied here), reducing the overall numbers of experimental animals needed to determine therapy efficacy, and ultimately improving our capacity to interpret the efficacy and robustness of experimental therapies under varied injury severity settings. For example, the use of AFS-defined subgroups can distinguish mice that will rapidly recover spontaneously and have small, constrained lesions (AFS>2) from those mice that will undergo very little spontaneous recovery because of larger, near full-width lesions (AFS<1) with essentially 100% accuracy. Thus, if experimental therapies are applied on or after 3 days, any improvement in recovery resulting in the return of stepping function across a high proportion of the AFS<1 cohort is more likely to reflect a meaningful treatment effect rather than spontaneous recovery or any procedural bias inadvertently created during the injury surgery. Notably, if the same experimental therapy is applied to AFS>2 mice, it is unlikely that any positive treatment effect would be identified, since natural functional recovery is very rapid and lesions are already constrained in size with extensive astrocyte bridging and preserved axonal tracts. However, any decrease in function or exacerbation in lesion size associated with treatment in these AFS>2 mice may suggest a deleterious effect of treatment on natural wound healing, which is equally important to identify. Information on both positive and negative treatment effects leading to a conversion or regression in the recovery class is readily incorporated into the confusion matrix and cosine similarity evaluation approach outlined in this work, enabling a robust evaluation of new treatments. We are actively exploring experimental interventions that are surgically delivered locally during the sub-acute injury phase on or after 3 days post injury to modulate endogenous wound repair processes and using this analytical framework to decipher treatment effects.

The analytical framework presented here for predicting recovery trajectories is not without limitations that should be addressed and improved upon in future work. For instance, the current framework only incorporates OF behavioral data from two timepoints (d1 and d3) to establish subgroup classifications. Given the underlying complexity and inherent variability in spontaneous animal behavior and its observation, there is high potential for misclassification using only acute OF data, which could undermine the robustness of predictions. Although our current approach performed very well with this limited input data, future work should extend the way the subgroup classification is performed to strengthen the assessment of injury severity acutely, such as by incorporating early kinematics evaluations. We could also consider obtaining other types of information about lesion severity ideally using minimally invasive methods already employed clinically, such as identifying the extent of intraparenchymal hemorrhage by magnetic resonance imaging (MRI) or ultrasound (US) imaging^34^, or characterizing neural tissue loss by non-invasive blood testing and quantitative immunoassays to measure serum biomarkers like GFAP or neurofilament (NF-L)^35^. The current predictive trajectory model has only been validated on a single type of injury, requires that administration of experimental treatments be delayed to, or after, 3 days post injury, and was used to make predictive evaluations only up to 2 weeks post injury. These constraints were selected because they aligned with the specific experimental paradigm we are actively pursuing in the lab and because we saw no meaningful increase in recovery by OF assessments beyond 2 weeks. It will be essential to have enhanced temporal flexibility for treatment administration and a capacity to account for treatment-dependent recovery effects that may take longer than two weeks to begin to manifest. It will be important to extend the analytical framework to other injury types (e.g. contusion injuries) and integrate age at the time of injury as an experimental variable, since these parameters can significantly impact recovery outcomes^2^. The decision to limit ourselves to 3 subgroups was based on our extensive observations with this partial crush injury model. However, as we accrue data from more experimental animals and potentially incorporate additional readouts into the AFS metric, it may be possible to expand the number of classified subgroups beyond the current framework to further enhance the accuracy of the recovery trajectories that are predicted. Powerful machine learning approaches including supervised and deep learning methods have begun to be incorporated into predictive recovery modeling after stroke and traumatic brain injury. These models are capable of integrating a variety of acutely derived features taken from electrophysiology, imaging, or behavioral evaluations^36^. There are opportunities to leverage these new tools and techniques to integrate multimodal acute injury data (kinematics, neuroimaging, serum markers, and/or electrophysiology) and build even more powerful predictive models of recovery in the future.

In conclusion, this work describes the development of an analytical framework to predict the trajectory of natural functional recovery and injury severity in mice subjected to a standardized partial crush SCI. The validation of this analytical framework outlined here supports its incorporation into the evaluation of experimental treatments administered in the sub-acute injury phase of SCI to neutralize procedural bias, minimize animal numbers, and provide a probabilistic basis for evaluating whether interventions enhance or suppress wound repair processes.

## Materials and Methods

### Surgical procedures

All surgical procedures were approved by BU IACUC (PROTO202100013) and performed in a designated surgical facility. All studies were performed on female and male C57BL/6 mice purchased from The Jackson Laboratory (JAX Cat#000664). Mice were 8-14 weeks of age at the time of spinal cord injury onset. All surgical procedures were conducted under general anesthesia using continuous isoflurane (0.5-2.0%) in oxygen-enriched air. Immediately following induction, subcutaneous Buprenorphine was administered as an analgesic. Animals were fitted into a stereotaxic frame with their heads secured with ear bars (David Kopf, Tujunga, CA) and their noses placed into a purpose-built anesthesia delivery and ventilation system. Eye ointment was applied before surgery.

To perform the laminectomy surgery and spinal cord forcep partial crush injury, using the aid of a stereo microscope, a 10mm skin incision was made oriented parallel to the spine. Subcutaneous fat at the incision site was removed and two small incisions in the muscle on both sides of the vertebrae at the intended laminectomy site were made using small spring scissors. Two small incisions in the lamina of the T10 vertebrae were made and the excised spinous process was removed using Dumont No.2 SP forceps. The forcep crush was applied using McPherson Tying Forceps, 4” Straight Ophthalmic (Premium Instruments), with a grommet spacer of 0.4 mm width and a tip width of 0.5 mm to compress the entire spinal cord laterally from both sides for 5 seconds. After crush injury the muscle incisions were sutured closed using 6-0 vicryl degradable sutures and the skin was closed using 6-0 prolene sutures in a subcuticular pattern. Subcutaneous Buprenorphine was administered every 12 hours for at least 72 hours post-surgery.

Mice receiving a saline or coacervate vehicle treatment received revision surgeries at 3 days post-injury. Mice were fitted into the stereotaxic frame as described above. Saline or coacervates were loaded into pulled borosilicate glass micropipettes (WPI #1B100-4) that were ground to a 35° beveled tip with 150–250 μm inner diameter after formulation with sterile-filtered components. Glass micropipettes were mounted to the stereotaxic frame and high-pressure polyetheretherketone tubing and connectors were used to attach micropipettes to a 10 μL syringe (Hamilton, Reno, NV, #801 RN) that was mounted into a syringe pump (Pump 11 Elite, Harvard Apparatus). The skin was reopened by clipping prolene sutures with spring scissors and the surgical site was re-exposed by gently peeling away cut tissue after removing vicryl sutures. Saline (1xPBS) mice received 4 separate injections made according to the following coordinates from the midline of the injury epicenter: +/- 0.4 mm L/M and +/- 0.2 A/P and -0.7 mm D/V. Each injection was 0.35 µl in volume, injected at a rate of 0.2 µl/min. BDA (10 kDa—Thermofisher, #D1956) loaded coacervates were prepared as previously described^31^. Coacervate injections were performed bilaterally 0.4 mm from either side of midline of the spinal cord at the lesion epicenter and 0.7 mm from the dorsal surface of the spinal cord. Each injection was 0.4 µl in volume, injected at a rate of 0.2 µl/min. Once the complete volume of saline solution or coacervate had been injected the micropipette was allowed to dwell in the spinal cord at the injection site for an additional 3 minutes before being slowly removed from the spinal cord incrementally over a 1-min period. After injection, the muscle incisions were sutured closed using 6-0 vicryl degradable sutures and the skin was closed using 6-0 prolene sutures in a subcuticular pattern. Subcutaneous Buprenorphine was administered every 12 hours for at least 72 hours post-surgery.

### Behavioral testing

Open field (OF) locomotion assessments were performed using a discrete 6-point scale that has been used extensively before^24^ and is based on the mouse’s ability to perform voluntary hindlimb movements, stepping, and/or weightbearing (range of scoring spans 0-5: 0 indicating no movement, voluntary or involuntary, and 5 indicating stepping and weightbearing with no disability). The left and right hindlimb were scored separately and averaged in all representations used here. For all testing, 2 observers simultaneously, but independently, scored the locomotion of a single mouse. OF scoring was performed 1, 3, 7, 10, and 14 days post-injury. Taking the area under the curve (AUC) of the OF score trace from days 1 to 14 (AUC OF_d1-14_) was used to generate a single relative recovery metric.

Grid walk (GW) testing was performed over a 3-minute test period using a custom-made, enclosed apparatus consisting of a metal grid (bar width = 1 mm, grid spacing = 1/2” x 1/2”, entire grid = 16”X10”) and Clear Perspex siding. The testing was recorded using a video camera (Sony - HDRCX405 HD) in MP4 format at 30 frames per second. GW testing was performed 7-, 10-, and 14-days post-injury. The left and right hindlimb stepping activity was evaluated separately and characterized into 3 distinct categories: plantar placement, slight slip, and total slip. Plantar placement designation required the mouse to place the designated hindlimb on the grid bar and bear weight without slipping. Slight slip characterization was indicated when a mouse was able to place their hindlimbs successfully on the grid, but the hindlimbs slipped off the grid once any weight was applied. Total slip was characterized by the mouse failing to place the stepping hindlimb onto the grid, resulting in the hindlimb falling through the spacing in the grid. To calculate a GW metric, we assigned successful plantar placed steps with a weighting of 4 points, slight slips a weighting of 2 points, and total missed steps through the grid a weighting of 1 point. The weighted average assessment of stepping quality was computed by summing the multiplication of the category weighting and the percentage of steps assigned to that category, with mice achieving perfect, 100% plantar placement receiving a weighted average score of 4. If the mouse was incapable of executing any stepping behavior during the 3-minute testing period, it received a score of 0 points on the weighted-average metric. To integrate both the quality of stepping events (the weighted average) and the level of activity during the testing periods, we generated a Composite Grid Walk score, which involved multiplying the weighted average by the square root of the total number of observed plantar placement stepping events. Taking the AUC for the Composite Grid Walk score from day 7 to 14 was used to generate a single outcome recovery metric for the GW test (AUC GW_d7-14_).

### Kinematic Data Acquisition and Analysis

#### Video Acquisition

Mice were habituated on a motorized treadmill (#761181, Data Sciences International) a few days prior to experimentation with an increasing belt speed from 3 to 15 cm/s. Locomotor behavior was recorded using a GoPro Hero 6 camera at 240 frames per second while mice walked on the treadmill. Treadmill speed was maintained at 7cm/s for SCI and healthy mice. All videos were imported into Shotcut (v 25.07.26) for preprocessing. Videos were trimmed into single continuous walking bouts, cropped to remove background and non-relevant areas of the arena, and exported in an mp4 format to be processed on DeepLabCut (DLC).

#### Pose Estimation with DeepLabCut

Markerless pose tracking was performed using DeepLabCut (DLC: v2.3.9). A set of anatomical landmarks (iliac crest, hip, knee, ankle, metatarsophalangeal (mtp) joint, and toe) were manually labeled on a subset of frames from multiple animals at varying speeds and sessions. The labeled dataset was then used to train the neural network in a 90/10 train-test split. Network training was performed on Boston University’s Shared Computing Cluster (SCC) using a ResNet-152 backbone with Imgaug-based image augmentation. The network was trained for 700,000 iterations, yielding a final training error of 2.37 and test error of 7.93, consistent with high-quality tracking for high-speed mouse locomotion video. A p-cutoff of 0.06 was applied to filter low-confidence predictions.

#### Automated Kinematic Extraction (ALMA)

Pose-estimation outputs were processed using the open source Automated Limb Motion Analysis (ALMA) v2.0 toolbox^28^. ALMA automatically extracted limb kinematic parameters including stride length (cm), drag duration (s), cycle duration (s), stance and swing percentages, and other metrics related to the animal’s locomotion. All default ALMA filtering and stride-detection parameters were used unless otherwise specified.

### Transcardial perfusion and immunohistochemistry

At 3 or 14 days post SCI, mice underwent terminal anesthesia by overdose of isoflurane and were then perfused transcardially with heparinized saline and 4% paraformaldehyde (PFA) using a peristaltic pump at a rate of 7 ml/min. Approximately 10 ml of heparinized saline and 50 ml of 4% PFA were used per animal. Spinal columns were dissected and post-fixed in 4% PFA for 6–8 h followed by cryoprotection in 30% sucrose with 0.01% sodium azide in Tris-Buffered Saline (TBS) for at least 3 days at 4 °C prior to cryosectioning. Spinal cords were cut in the horizontal plane into sections (30 μm thick) using a cryostat (Leica CM1950 Cryostat). Tissue sections were stored in 96-well plates immersed in 1X TBS and 0.01 % sodium azide solution at 4 °C. For immunohistochemistry (IHC), tissue sections were stained using a free-floating staining protocol outlined in detail previously^7,30^. Following 1N HCl antigen retrieval and serial washes, the tissue sections were blocked and permeabilized using 1× TBS containing 5% Normal donkey serum (NDS, Jackson ImmunoResearch Laboratories) and 0.5% Triton x-100 for 1 h. Tissue sections were then stained with primary antibodies overnight in 1× TBS containing 0.5 % Triton X-100. The following primary antibodies were used in this study: anti-Rat Gfap (Invitrogen, 13–0300, 1:1000); anti-Guinea Pig Gfap (Synaptic Systems, 173 308, 1:500); anti-Goat Cd13 (R&D systems, AF2335, 1:500), anti-Rabbit 5-HT (Serotonin) (Immunostar, #20080, 1:1000); anti- Rat MBP (Sigma, MAB386, 1:500). Tissue sections were washed three times and incubated with appropriate secondary antibodies diluted at 1:250 in 1X TBS containing 0.5 % Triton X-100 and 5 % NDS for 2 h. All secondary antibodies were affinity purified whole IgG (H + L) purchased from Jackson ImmunoResearch Laboratories, with donkey host and target specified by the primary antibody. Cell nuclei were counter stained with 4′,6′-diamidino-2-phenylindole dihydrochloride (DAPI; 2 ng/ml; Molecular Probes) prior to mounting onto glass slides and then cover slipped with ProLong Gold (Invitrogen, Cat# P36934) anti-fade reagent. Stained sections were imaged using epifluorescence and deconvolution epifluorescence microscopy on an Olympus IX83 microscope.

### Subgroup classification and behavioral recovery trajectory analysis

We used a latent class growth analysis and growth mixture modeling approach to separate the population data from the behavioral evaluations into distinct subgroups (latent classes) that follow different longitudinal trajectories. Subgroup classifications were performed by taking the AUC of the OF scores recorded from post injury days 1 to 3 (AUC OF_d1-3_). To simplify nomenclature, the AUC OF_d1-3_ parameter was designated the Acute Functional Score (AFS). The range of AFS values that were separated into the three distinct subgroup designations was determined by minimizing the root mean square error (RMSE) for fitted regressions applied to six unique, non-overlapping arrangements of three AFS subgroups across the 1-14 day interval of the OF data. A four-point parameter (4PL) sigmoidal regression was used in the fitting of the OF data in the following form:

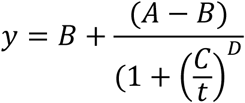

Where: A, B are the top and bottom plateaus respectively, C is the inflection point, and D is the Hill Slope.

All fitted regressions and goodness-of-fit parameters (RMSE, R^2^, Cumulative sum residuals (CUSUM), standard deviation of the residuals (Sy.x)) were generated using OF data from days 1, 3, 7, 10, and 14 and performed using Prism 10 (GraphPad Software Inc, San Diego, CA). Three unique 4PL sigmoidal regressions designated OF regressions 1, 2, and 3 were generated by fitting the data separated into the three unique AFS-defined subgroups and including all animals assigned to those subgroups.

To evaluate the suitability of an individual animal being assigned to any one of the three subgroups based on OF trajectory regressions, we used leave-one-out cross-validation (LOOCV) and computed posterior model probabilities (PMP).

To calculate PMP values for a given animal, we first assessed the residuals (r_t_) at each timepoint associated with fitting an individual animal’s data to each of the three OF trajectory regressions making sure to leave out that animal from any of the datasets used to define the regressions.

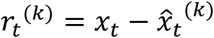

Where:

*x*_*t*_ is observed value at time t
*x̂_t_* is predicted value from the OF trajectory regression
where k = 1, 2, or 3 for OF trajectory regression 1, 2 or 3.

Residual Sum of Squares (RSS) were then computed across the 1, 3, 7, 10, and 14 day timepoints by performing the following calculation:

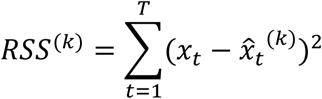

Using the RSS values and the Sy.x for each OF trajectory regression, we then calculated the relative likelihood for each regression using the following equation:

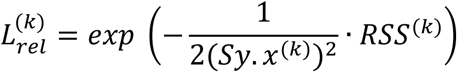

Then, assuming equal prior probability, the posterior model probabilities (PMP) were then evaluated by comparing the likelihoods of the three regressions using the following equation:

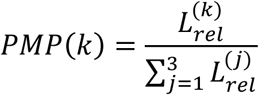

GW composite scores across the three subgroups were fitted with a simple linear regression using Prism 10 (GraphPad Software Inc, San Diego, CA). To evaluate the suitability of an individual animal being assigned to any one of the three subgroups based on GW trajectory linear regressions, we used leave-one-out cross-validation (LOOCV) and computed posterior model probabilities (PMP) using the same analytical methods described above for the OF analysis.

Since OF and GW data were strongly correlated, and therefore not independent, we computed a combined evaluation metric by averaging the OF and GW PMPs. We referred to this parameter as combined PMP. Recovery classes were assigned according to the highest PMP for the associated trajectory for the OF and GW evaluations or the highest average PMP for the combined evaluation. Confusion matrices were generated by calculating the percentage of mice from the observed AFS subgroup displaying Class 1, 2, and 3 type recovery. In these confusion matrices, the sum of the columns is equal to 100%. Cosine similarity (CS) analysis was performed by flattening a given confusion matrix into a 9-element vector, calculating the dot product between it and the ideal identity matrix, and then dividing by the multiplication of the two vector magnitudes.

### Quantification of IHC

Immunohistochemical staining intensity quantification for the various antibodies was performed on tiled images prepared on the Olympus IX83 epifluorescence microscope, acquired at a standardized exposure time using a 10x objective and the raw/uncorrected intensity settings. Quantification of total antibody staining in tissue was performed using NIH Image J (1.51) software and the plot profile plugin as we have done before^4^. Total values for IHC staining intensities were determined by taking the integral (area under the curve) of the intensity profile. Lesion size was determined by outlining Gfap-negative regions of tissue and measuring area. Astrocyte bridging was measured by taking line profiles across the spinal cord laterally and summing total Gfap measuring Gfap-positive staining across the cord as a percent of total cord width.

### Statistical analysis

Statistical evaluations were conducted by one-way or two-way ANOVA with post hoc independent pairwise analysis via Tukey’s or Sidak’s multiple comparisons test where appropriate using Prism 10 (GraphPad Software Inc, San Diego, CA). Correlation analyses were performed using Prism 10 with statistical evaluations conducted using t-test for Pearson’s linear correlation. Across all statistical tests, significance was defined as p-value <0.05. All graphs show mean values plus or minus standard error of mean (s.e.m.) as well as individual values overlaid as dot plots.

## Acknowledgements

We thank the Micro and Nano Imaging (MNI) core in the Biomedical Engineering Core Facilities at Boston University for microscope support. This research was supported by funding from: Boston University Start-Up Funds (T.M.O.), Boston University’s Undergraduate Research Opportunities Program (UROP) (H.P.), Boston University BUNano PhD Fellowship (P.S.W.), Paralyzed Veterans of America Research Foundation (T.M.O.), Wings for Life (T.M.O.), Ruth L. Kirschstein Predoctoral Individual National Research Service Award NIH NINDS (F31 NS145754 to L.F.H), and Maximizing Investigators’ Research Award (MIRA) NIH NIGMS (R35GM154942 to T.M.O.) (the content in this manuscript is solely the responsibility of the authors and does not reflect the official views of the NIH).

## Author contributions

T.M.O. conceptualized and led the overall project. T.M.O. designed, guided, and supervised all experiments. T.M.O., K.L., and L.F.H. performed surgeries. T.M.O., K.L., L.F.H., H.P., G.C.O., and P.S.W. conducted behavioral testing and analysis. G.C.O. conducted kinematics assessments. K.L. and L.F.H. conducted histological processing and microscope imaging. T.M.O., K.L., L.F.H, H.P., G.C.O., and P.S.W. analyzed data. T.M.O. wrote the paper with input and edits from all co-authors.

## Data and materials availability

All data needed to evaluate the conclusions in the paper are present in the paper and/or the Supplementary Materials. The data that support the findings of this study are available from the corresponding author upon reasonable request.

