## Supplementary Information for "Predicting recovery trajectories and injury severity following partial crush spinal cord injury in mice"

Four-point parameter (4PL) sigmoidal curve fitting of OF testing data

$$y = B + \frac{(A - B)}{\left(1 + \left(\frac{C}{t}\right)^{D}\right.}$$

| | | Population | AFS<1 | AFS1-2 | AFS>2 |
|---|---|---|---|---|---|
| **Fitted Parameters for Equation** | A | 2.90 | 1.34 | 3.34 | 4.67 |
| | B | 0.39 | 0.06 | 0.35 | 0.52 |
| | C | 7.04 | 8.47 | 5.14 | 4.20 |
| | D | 2.02 | 5.63 | 2.91 | 1.12 |
| **Goodness of Fit** | $R^2$ | 0.27 | 0.31 | 0.68 | 0.72 |
| | Sy.x | 1.26 | 0.75 | 0.80 | 0.62 |
| | RMSE | 1.25 | 0.74 | 0.78 | 0.60 |
| | AICc | 115.10 | -61.88 | -19.46 | -50.48 |

| **Calculated Days to OF score using Equation** | | | | |
|---|---|---|---|---|
| **OF Score** | **Population** | **AFS<1** | **AFS1-2** | **AFS>2** |
| **1** | 4.00 | 10.14 | 3.31 | 0.69 |
| **2** | 9.37 | - | 5.52 | 2.49 |
| **3** | - | - | 10.40 | 5.98 |
| **4** | - | - | - | 18.24 |

**Figure S1:** Four-point parameter (4PL) sigmoidal regression fits for the population data set (all mice in study, n=48) and the three subgroups for the open field (OF) behavioral testing. The mean plateau OF score is derived from the value for B for the 4PL equation fit and the mean timepoint of recovery acceleration (inflection point) is derived from parameter C for the 4PL equation fit. $R^2$ values indicate the fit for the 4PL equation factoring in the variability across all samples within the group. For the population and all subgroups, the $R^2$ for the regression fit to the mean values of the OF scores recorded from day 1 to 14 post injury was ~0.99. For each of the fitted 4PL regressions for the population and subgroups, the times taken to reach OF scores of 1, 2, 3, and 4 was also calculated.

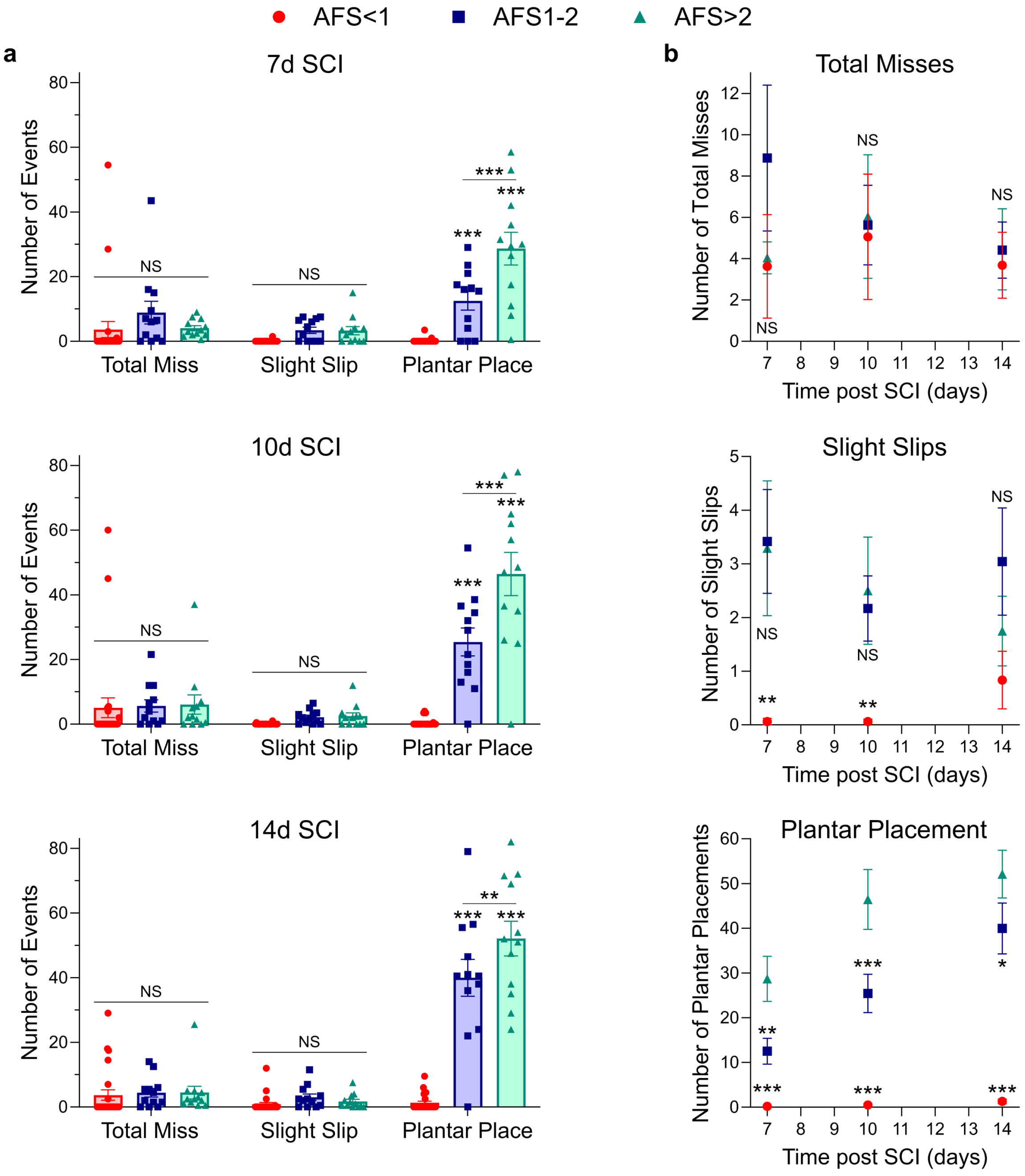


**Figure S2:** Counts for the number of events for the individual Grid Walk (GW) testing parameters (total miss, slight slip, plantar placement) according to AFS subgroup designations and separated by **a.** time post SCI (7d, 10d, 14d) or **b.** parameter. *** p-value<0.0001, ** p-value<0.005, * p-value<0.05, Not significant (NS), Two-way ANOVA with Tukey's multiple comparisons test. In **a.** comparisons are referencing the AFS<1 group for the same parameter except where indicated. In **b.** all comparisons are made to the AFS>2 at the same timepoint. Graphs in a&b. show mean + standard error of mean (s.e.m.) with individual data points overlayed in a.

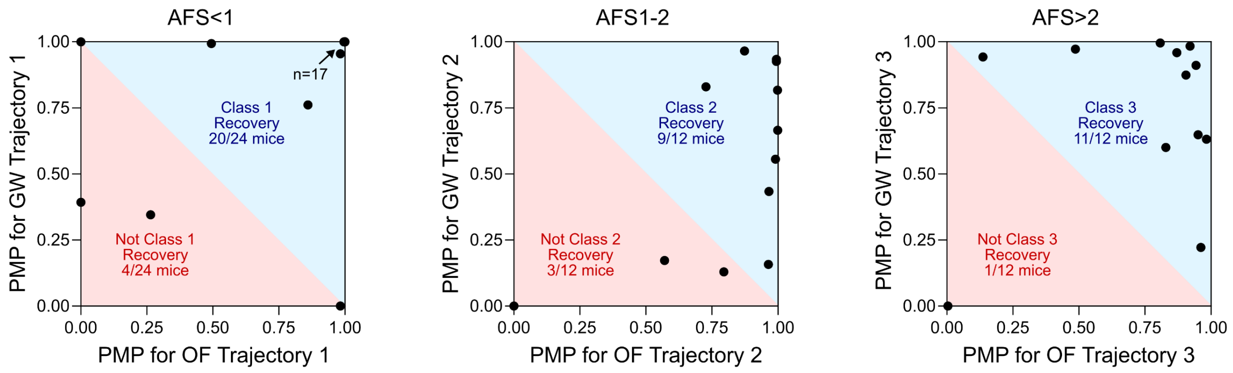


**Figure S3:** Comparisons of PMP values for GW and OF trajectories associated with the three subgroups. The blue shaded regions provide a graphical representation of whether the Combined PMP value is greater than 0.5 for the associated subgroup. Mice falling into the blue shaded region on their AFS defined subgroup are considered to have recovery trajectories consistent with that AFS defined subgroup as defined by the combined PMP metric. By contrast, mice falling into the red shaded region have converted to a higher or lower class of recovery according to the combined PMP metric.

a

**PCA Factor Loadings**

| Parameter | PC1 | PC2 |
|---|---|---|
| **ankle joint flexion (deg)** | **0.88** | 0.09 |
| **mtp joint flexion (deg)** | **0.86** | 0.17 |
| mean toe-to-crest distance (cm) | 0.71 | 0.06 |
| min toe-to-crest distance (cm) | 0.71 | 0.01 |
| max toe-to-crest distance (cm) | 0.70 | 0.11 |
| drag percentage (%) | 0.70 | 0.21 |
| hip joint flexion (deg) | 0.68 | 0.32 |
| swing percentage (%) | 0.66 | -0.34 |
| drag duration (s) | 0.63 | -0.03 |
| swing duration (s) | 0.58 | -0.28 |
| ankle joint extension (deg) | 0.57 | 0.69 |
| mtp joint extension (deg) | 0.55 | -0.04 |
| cycle duration (no. frames) | 0.45 | -0.27 |
| hip joint extension (deg) | 0.39 | 0.54 |
| stride length (cm) | 0.19 | 0.33 |
| knee joint flexion (deg) | -0.04 | 0.68 |
| **toe-to-crest distance SD (cm)** | -0.12 | **0.80** |
| step height (cm) | -0.30 | -0.56 |
| cycle velocity (cm/s) | -0.35 | 0.63 |
| max velocity during swing (cm/s) | -0.37 | 0.47 |
| knee joint SD (deg) | -0.44 | -0.66 |
| knee joint amplitude (deg) | -0.45 | -0.65 |
| knee joint extension (deg) | -0.48 | -0.06 |
| stance duration (s) | -0.51 | 0.06 |
| hip joint amplitude (deg) | -0.58 | 0.33 |
| ankle joint amplitude (deg) | -0.60 | 0.60 |
| ankle joint SD (deg) | -0.63 | 0.61 |
| hip joint SD (deg) | -0.64 | 0.34 |
| mtp joint SD (deg) | -0.65 | -0.27 |
| stance percentage (%) | -0.66 | 0.34 |
| mtp joint amplitude (deg) | -0.68 | -0.20 |

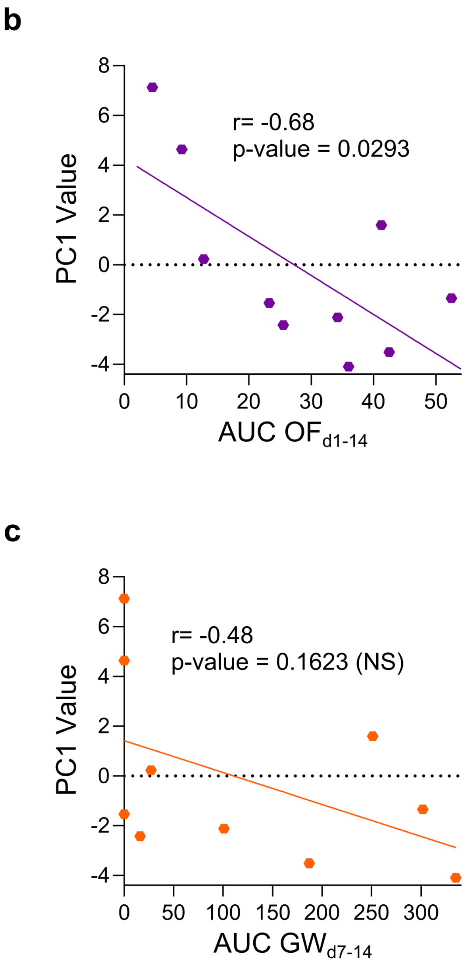


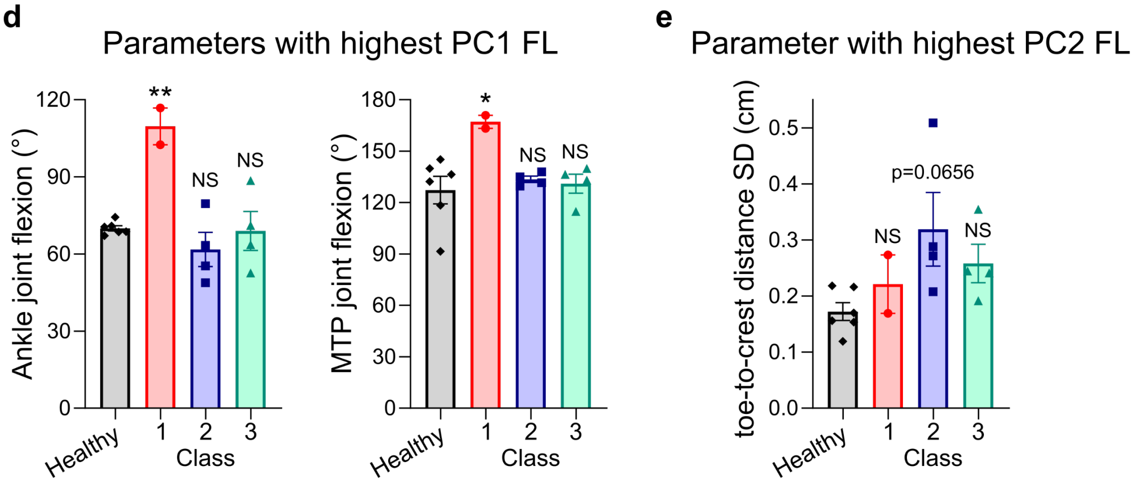


**Figure S4:** **a.** List of all kinematics parameters used in the PCA analysis and their corresponding PC1 and PC2 factor loadings (FL). **b.&c.** Correlation analysis of PC1 values and OF (b) and GW (c) scores, with the PC1 values showing significant correlation with OF (r=-0.68) but not GW (r=-0.48) results, p-value indicated on graph, t-test for Pearson's linear correlation. **d.&e.** Results for kinematics parameters that had FL>0.8 for PC1 (d) and PC2 (e). ** p-value<0.005, * p-value<0.05, Not significant (NS), One-way ANOVA with Tukey's multiple comparisons test. Graphs in d&e. show mean + standard error of mean (s.e.m.) with individual data points overlayed.

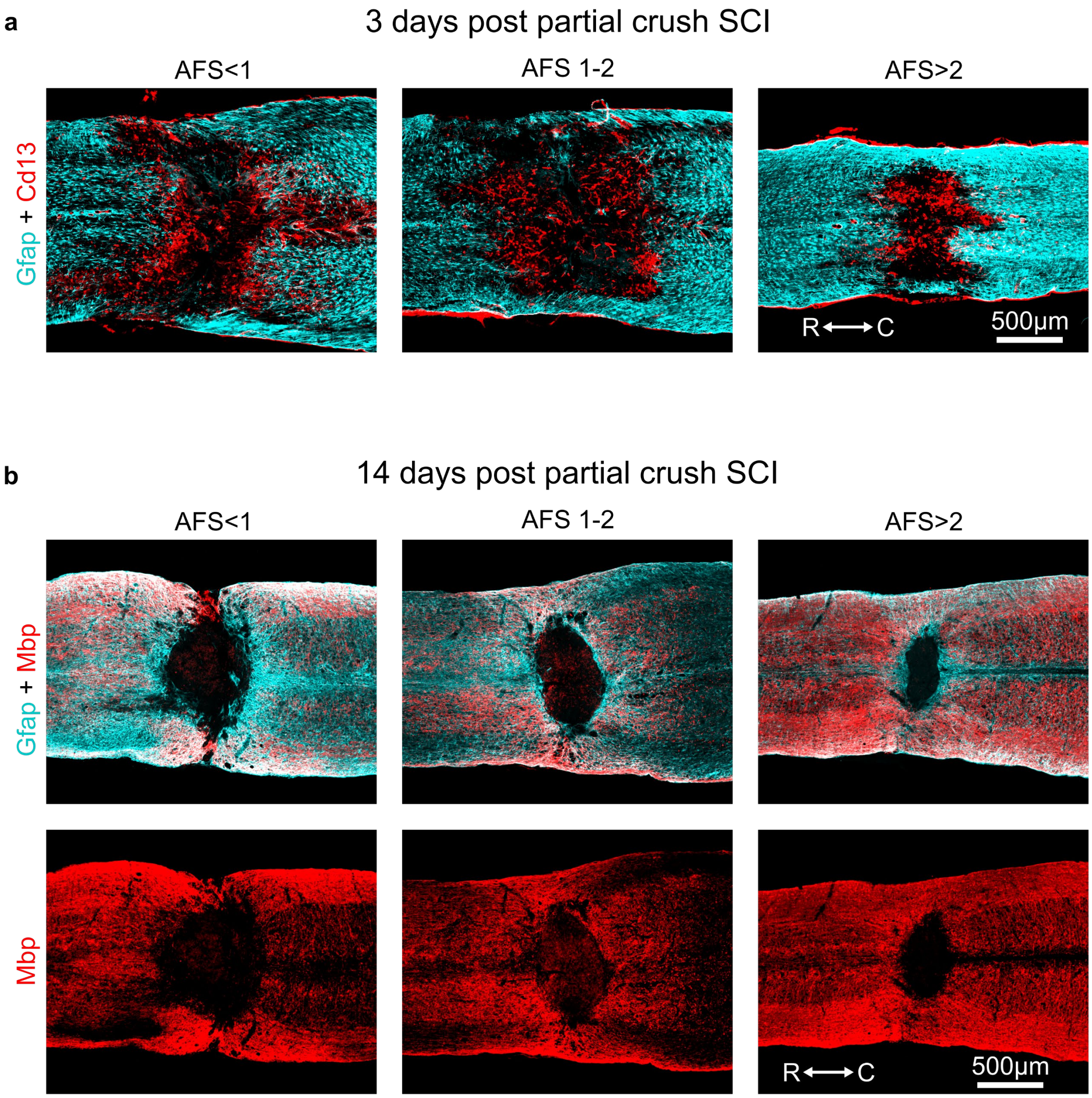


**Figure S5:** **a.** Representative survey IHC images of SCI lesions from the three AFS defined subgroups at 3 days after partial crush injury showing Gfap-positive astrocytes and Cd13-positive immune and stromal cells within the lesion cores. **b.** Representative survey IHC images of SCI lesions from the three AFS defined subgroups at 14 days after partial crush injury showing regions of myelin basic protein (Mbp) loss within the lesion core. Intact myelin basic protein (Mbp) positive tissue at the lateral margins of the tissue surrounding the lesion cores suggests that Gfap-positive astrocyte bridging is likely derived from spared surviving neural tissue.
